# Fgf3 is crucial for the generation of monoaminergic cerebrospinal fluid contacting cells in zebrafish

**DOI:** 10.1101/477323

**Authors:** Isabel Reuter, Jana Jäckels, Susanne Kneitz, Jochen Kuper, Klaus-Peter Lesch, Christina Lillesaar

**Affiliations:** Division of Molecular Psychiatry, Center of Mental Health, University of Würzburg, Germany; Department of Physiological Chemistry, Biocenter, Am Hubland, University of Würzburg, Germany; Structural Biology, Rudolf Virchow Center for Biomedical Research, University of Würzburg, Germany; Laboratory of Psychiatric Neurobiology, Institute of Molecular Medicine, I.M. Sechenov First Moscow State Medical University, Moscow, Russia; Department of Neuroscience, School for Mental Health and Neuroscience (MHeNS), Maastricht University, Maastricht, The Netherlands; Department of Child and Adolescent Psychiatry, Psychosomatics and Psychotherapy, Center of Mental Health, University Hospital of Würzburg, Germany.

**Keywords:** Fgf-signalling, serotonin, dopamine, hypothalamus, central nervous system

## Abstract

In most vertebrates, including zebrafish, the hypothalamic serotonergic cerebrospinal fluid-contacting (CSF-c) cells constitute a prominent population. In contrast to the hindbrain serotonergic neurons, little is known about the development and function of these cells. Here, we identify Fibroblast growth factor (Fgf)3 as the main Fgf ligand controlling the ontogeny of serotonergic CSF-c cells. We show that *fgf3* positively regulates the number of serotonergic CSF-c cells, as well as a subset of dopaminergic and neuroendocrine cells in the posterior hypothalamus. Further, expression of the ETS-domain transcription factor *etv5b* is downregulated after *fgf3* impairment. Previous findings identified *etv5b* as critical for the proliferation of serotonergic progenitors in the hypothalamus, and therefore we now suggest that Fgf3 acts via *etv5b* during early development to ultimately control the number of mature serotonergic CSF-c cells. Moreover, our analysis of the developing hypothalamic transcriptome shows that the expression of *fgf3* is upregulated upon *fgf3* loss-of-function, suggesting activation of a self-compensatory mechanism. Together, these results highlight Fgf3 in a novel context as part of a signalling pathway of critical importance for hypothalamic development.

**Summary statement:** This study highlights Fgf3 in a novel context where it is being part of a signalling pathway of critical importance for development of hypothalamic monoaminergic cells in zebrafish.

## Introduction

Serotonin (5-hydroxytryptamine, 5-HT) is an ancient signalling molecule present in the nervous system of animals from cnidarian and bilaterian lineages (Hay-Schmidt, 2000; Kass-Simon and Pierobon, 2007; Moroz and Kohn, 2016; Stach, 2005). Accordingly, 5-HT modulates a variety of physiological processes and behaviours in most animals (e.g. Curran and Chalasani, 2012; Fossat et al., 2014; Gillette, 2006; Lucki, 1998; Trowbridge et al., 2011). In placental mammals, serotonergic neurons are, with a few potential exceptions (Ballion et al., 2002; Ugrumov et al., 1989), uniquely located in the raphe nuclei of the hindbrain (Deneris and Gaspar, 2018). In contrast, additional serotonergic cell populations are found in the forebrain and spinal cord in cartilaginous and bony fish, amphibians, reptiles, birds and monotremes (Lillesaar, 2011; Montgomery et al., 2016). Most of these non-raphe cell populations are truly serotonergic as they contain not only 5-HT, but also proteins required for 5-HT metabolism, packaging and transport. To which extent the development of different populations of serotonergic cells is controlled by the same paracrine signals and gene regulatory networks to ultimately express the mature serotonergic phenotype (i.e. capacity to synthesise 5-HT) is still unclear.

The ontogeny as well as the regulatory transcriptional networks of raphe serotonergic neurons are well described (Deneris and Gaspar, 2018; Deneris and Wyler, 2012; Flames and Hobert, 2011; Kiyasova and Gaspar, 2011; Lillesaar et al., 2007, 2009; McLean and Fetcho, 2004; Norton et al., 2005; Teraoka et al., 2004). In contrast, little is known about the development of the remaining populations in the central nervous system. In zebrafish, the hypothalamus contains by far the highest number of serotonergic cells (Lillesaar, 2011). These cells are small, bipolar cells with one thick process contacting the ventricle suggesting that they can communicate over longer distances via the cerebrospinal fluid (CSF), and the other process projecting locally in the brain (Lillesaar, 2011; Sano et al., 1983; Vígh et al., 2004). Anatomically, these cells are further separated into three clusters in teleost fish; the anterior, intermediate (i.) and posterior (p.) paraventricular organ clusters (Ekström et al., 1985; Kaslin and Panula, 2001), of which the latter is located around the posterior recess, a ventricular structure found in the hypothalamus of teleosts (Xavier et al., 2017). Overlapping expression of *tph1a*, *slc6a4b, ddc, mao* and *vmat2* (Anichtchik et al., 2006; Lillesaar et al., 2007; Norton et al., 2008; Sallinen et al., 2009; Yamamoto et al., 2011) shows an active 5-HT metabolism in the region. Notably, serotonergic and dopaminergic cells are largely intermingled populations in the hypothalamus (Filippi et al., 2009; Kaslin and Panula, 2001; McLean and Fetcho, 2004; Xavier et al., 2017; Yamamoto et al., 2010, 2011).

Progenitors giving rise to neuronal and glial precursors of the developing hypothalamus are located at the ventricular zone (Duncan et al., 2016; Xie and Dorsky, 2017). Eventually the progenitors exit the cell cycle, migrate laterally and continue differentiation. Lineage specific combinations of transcription factors control the fate of the precursors (Biran et al., 2015; Burbridge et al., 2016; Muthu et al., 2016; Ware et al., 2014; Xie and Dorsky, 2017). The precise set of paracrine molecules and transcription factors required in time and space for each cell type is still a matter of investigations, as is the extent of evolutionary conservation. In zebrafish, the location of the serotonergic cells in the posterior hypothalamus suggests that their fate is favoured by posteriorising signals such as Wnt and Fgf (Kapsimali et al., 2004; Xie and Dorsky, 2017), and inhibited by anteriorising signals such as late Shh expression (Mathieu et al., 2002; Muthu et al., 2016). Indeed, Fgf ligands, receptors and downstream targets are expressed in the posterior zebrafish hypothalamus (Bosco et al., 2013; Herzog et al., 2004; Jackman et al., 2004; Liu et al., 2013; Reifers et al., 1998; Topp et al., 2008). Similarly to raphe serotonergic neurons, hypothalamic serotonergic CSF-c cells along with dopaminergic cells depend on Fgf-signalling during development (Bosco et al., 2013; Koch et al., 2014; Teraoka et al., 2004). Furthermore, Etv5b, a member of the ETS-domain transcription factor family that is a direct downstream target of Fgf-signalling (Ornitz and Itoh, 2015; Raible and Brand, 2001; Roussigné and Blader, 2006), regulates the proliferation of serotonergic progenitors (Bosco et al., 2013).

Here, we are focusing on the hypothalamic serotonergic cells of zebrafish, a frequently used model organism in biomedical research. Using three different approaches for genetic manipulation, we identify Fgf3 as the main Fgf ligand critical for development of serotonergic as well as dopaminergic CSF-c cells and *arginine vasopressin (avp)*-expressing cells located in the posterior hypothalamus. Further, based on sequencing of the transcriptome of microdissected hypothalami we identify genes belonging to the Fgf-signalling pathway that are expressed in the developing hypothalamus, and demonstrate mild alterations of Fgf-signalling after impairment of Fgf3. With this information we acquire a better knowledge about the signalling networks promoting the ontogeny of central serotonergic cells.

## Material and Methods

### Fish husbandry and sample preparation

Two zebrafish (*Danio rerio*) strains were used for the experiments: AB/AB wildtypes and the *fgf3^t24152^ N*-ethyl-*N*-nitrosourea mutant in a Tü/Tü background *(lia*) (Herzog et al., 2004). Animal husbandry followed the animal welfare regulations of the District Government of Lower Franconia, Germany. Embryos, staged according to Kimmel at al. (1995) (Kimmel et al., 1995), were raised and maintained in Danieau’s solution (Cold Spring Harb. Protoc., 2011) with a 14/10-h light/dark cycle at 28°C. To prevent pigmentation 0.2 mM 1-phenyl-2-thiourea was added to Danieau’s solution after 24 hours post fertilisation (hpf). For immunohistochemistry or *in situ* hybridisation, embryos were dechorionated and fixed for 24 h at 4°C in 4% paraformaldehyde in phosphate buffered saline, subsequently dehydrated in increasing concentrations of methanol, and stored in 100% methanol at −20°C.

### Genotyping

Embryos with the *fgf3*^*t24152*^ allele were genotyped by PCR using HiDi SNP DNA Polymerase (Genaxxon Bioscience). The polymerase allowed for allele specific discrimination of the gene fragment containing the point mutation *fgf3*^*t24152*^ (G to A transition). Two PCR reactions were prepared to genotype each embryo. For the first reaction a forward (fwd) primer in which the last nucleotide at the 3’ end (G) was specific to the wildtype allele (fwd primer wildtype:

5’GCCAGTTCTAAAAGGCAGTG**G**3’), and for the second reaction a fwd primer in which the last nucleotide at the 3’ end (A) was specific to the mutant allele (fwd primer mutant: 5’GCCAGTTCTAAAAGGCAGTG**A**3’) were used, respectively. The reverse (rev) primer was identical for both reactions (rev primer: 5’TGCCGCTGACTCTCTCTAAG3’). Both PCR products were run on a 1.5% agarose gel. Depending on the embryo’s genotype, either a single band for the wildtype primer reaction (genotype: +/+), or a single band for the mutant primer reaction (genotype: -/-) was detectable. For the heterozygote genotype (+/-) a band was present in both reactions.

### *fgf3* impairment using morpholino and CRISPR/Cas9

For a morpholino based knock-down of *fgf3*, 0.5 mM of a splice-blocking morpholino (5’CCCGACGTGACATAACACTTACTGA3’, Gene Tools) targeting the splice donor site of exon 2/intron 2 of *fgf3* was injected into AB/AB fertilised eggs at the one-cell stage (Suppl. Fig. 1). Blocking the splicing of *fgf3* pre-mRNA lead to partial and complete inclusion of intron 2, which in both cases generated a nonsense amino acid sequence with a stop, thus resulting in a truncated Fgf3 protein. Efficiency of the splice morpholino was evaluated by reverse transcription (RT)-PCR using RevertAid First Strand cDNA Synthesis Kit (Thermo Fisher Scientific) and GoTaq polymerase (Promega) according to manufacturer’s instructions. The PCR products were separated on a 1.5% agarose gel, the bands were gel extracted using the GenElute Gel Extraction Kit (Sigma-Aldrich) and verified by Sanger sequencing (Eurofins).

**Fig. 1.**
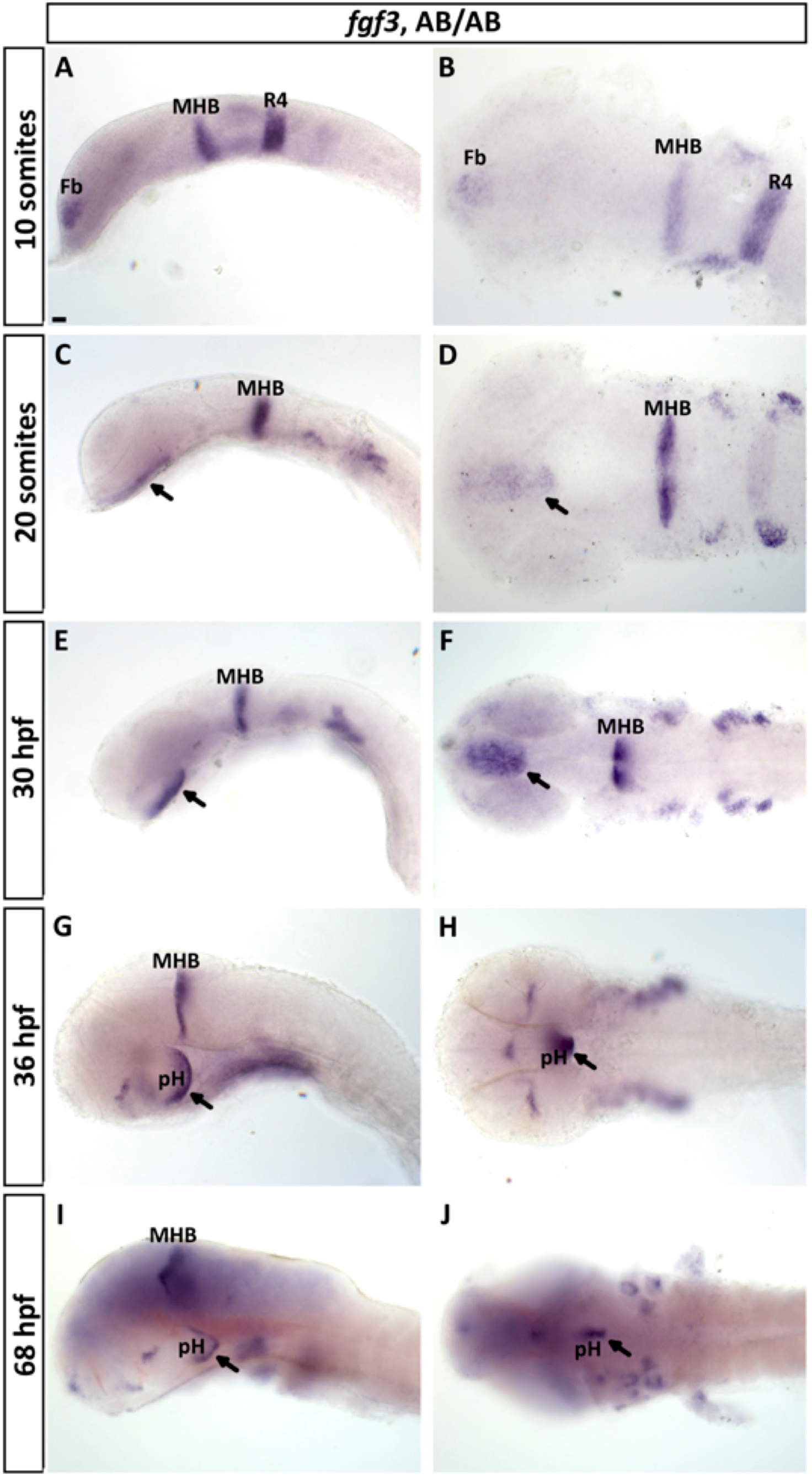
RNA *in situ* hybridisation reveals *fgf3* expression in the developing hypothalamus. **(A, B)** At 10 somites *fgf3* transcripts are detectable in the forebrain (Fb), at the mid-hindbrain boundary (MHB) and in rhombomere 4 (R4). **(C-F)** At 20 somites and 30 hpf *fgf3* expression is present in the hypothalamic primordium (arrow), and **(G-J)** at 36 and 68 hpf in the posterior hypothalamus (pH, arrow). At 68 hpf the expression is restricted to cells located at the ventricle. Left column depicts lateral views and right column ventral views of the embryos. Anterior to the left. Scale bar = 30 µm.

For the *fgf3* CRISPR/Cas9 strategy two different gRNAs targeting exon 1 and exon 2 of *fgf3* (Suppl. Fig. 2) were designed using the web tool CHOPCHOP (Labun et al., 2016; Montague et al., 2014). gRNA oligos (Suppl. Table 1) were annealed, cloned into vector DR274 (a kind gift from Keith Joung Addgene plasmid # 42250), *in vitro* transcribed using T7 RNA polymerase (a kind gift from Thomas Ziegenhals and Utz Fisher) and purified by Roti-Aqua-phenol/chloroform/isoamylalcohol (Roth) extraction. AB/AB embryos were co-injected at the one-cell stage with a cocktail containing both gRNAs (100-125 ng/µl each) and Cas9-NLS protein (300 ng/µl, *S. pyogenes*, New England Biolabs). To verify that the gRNAs induced indel mutations at the expected sites of *fgf3*, the target sites were amplified by PCR using fwd and rev primers listed in Suppl. Table 1 followed by separation of the PCR products on a 3% high-resolution NuSieve 3:1 agarose gel (Lonza), PCR clean up using the GenElute PCR Clean-up Kit (Sigma-Aldrich) and Sanger sequencing (Eurofins).

**Fig. 2.**
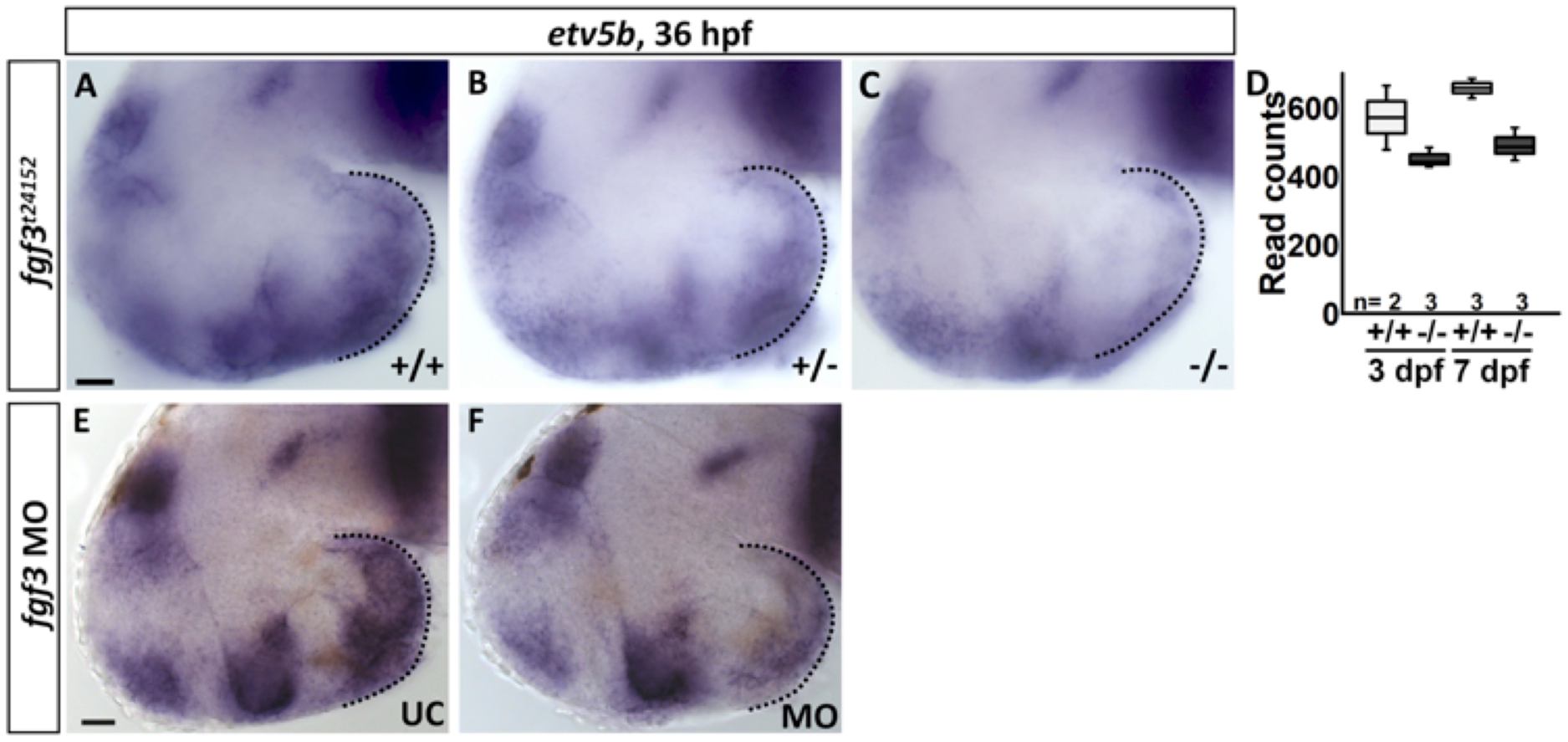
*fgf3*^*t24152*^ mutants and *fgf3* morphants exhibit a reduced expression of *etv5b* in the posterior hypothalamus. **(A-C, E, F)** Light microscopic pictures of wildtype (+/+), heterozygous (+/-) and homozygous (-/-) *fgf3*^*t24152*^ mutant siblings, and uninjected control (UC) and *fgf3* morphant (MO) siblings processed for RNA *in situ* hybridisation for *etv5b* at 36 hpf. Dotted line indicates posterior edge of hypothalamus where *etv5b* expression is present. Notably, the hypothalamic *etv5b* expression is weaker in +/-than in +/+ embryos, and strongly reduced in -/- embryos. Similarly, *etv5b* expression is weaker in MO embryos than in UC embryos. Lateral views, anterior to the left. Scale bar = 30 µm. **(D)** Read counts of *etv5b* obtained by RNA sequencing of dissected hypothalami of +/+ and *fgf3*^*t24152*^ -/- mutants at 3 and 7 dpf. Boxplots show median, 25-75% percentile, min/max whiskers. n = number of analysed replicates.

For both, morpholino and CRISPR/Cas9, strategies uninjected stage matched siblings were used as controls. Additionally, Cas9-NLS protein only injected embryos were used as injection controls for the CRISPR/Cas9 experiments.

### Morpholino toxicity assay using acridine orange

Live 24 hpf *fgf3* morpholino injected embryos and uninjected wildtype controls were incubated in 5 µg/ml acridine orange dissolved in Danieau’s solution for 30 min at 28°C, rinsed and imaged to reveal cell death.

### Whole mount immunohistochemistry

Whole mount immunohistochemistry was performed on embryos with heads dissected free of skin, eyes and jaws for better antibody penetration. Dissected embryos were incubated in immuno blocking buffer (10% normal sheep serum, 0.2% bovine serum albumin in phosphate buffered saline with 0.1% Tween 20 (PBT)) for 2 h at room temperature. Embryos were subsequently labelled with mouse anti-TH1 (1:500; MAB318, Millipore) and rabbit anti-5-HT (1:2500; S5545, Sigma-Aldrich) diluted in blocking buffer for 3 days at 4°C with gentle shaking. After washes in PBT, to reveal 5-HT and TH1 immunoreactivity the specimens were incubated for 2 days at 4°C in secondary antibodies conjugated with Alexa Fluor 488 (goat anti-rabbit, 1:1000; A-11034, Thermo Fisher Scientific) and Alexa Fluor 568 (donkey anti-mouse, 1:1000; A-10037, Thermo Fisher Scientific) diluted in immuno blocking buffer.

### Whole mount RNA *in situ* hybridisation

To detect mRNA transcripts whole mount *in situ* hybridisation was performed on fixed embryos as described elsewhere (Thisse and Thisse, 2008). In brief, *in vitro* transcribed digoxygenin (DIG) labelled antisense RNA probes (Suppl. Table 2) were synthesised using linearised plasmid template DNA, suitable RNA polymerases and DIG RNA labelling mix (Roche) following manufacturer’s recommendations. Prehybridisation, hybridisation and stringency washes were performed at 65°C. Alkaline phosphatase conjugated to anti-DIG Fab fragments (1:5000, Roche) was used to label hybridised transcripts, and to enable subsequent colour precipitation with NBT/BCIP solution (Roche).

### Sample preparation and data analysis of RNA sequencing

Hypothalami of 3 and 7 dpf homozygous wildtype embryos (Tü/Tü) and *fgf3*^*t24152*^ homozygous mutant embryos (in Tü/Tü background) were dissected (Fig. 11A). Wildtype embryos were generated by crossing previously identified homozygous wildtype adult fish, which were siblings to the heterozygous parents used to generate *fgf3*^*t24152*^ homozygous mutant embryos. Thus, the homozygous wildtype and the *fgf3*^*t24152*^ homozygous mutant embryos used for the RNA sequencing analysis were cousins. The *fgf3*^*t24152*^ homozygous mutant embryos were identified by their characteristic fused otolith phenotype (Herzog et al., 2004). Dissections were performed on ice in a petri dish containing ice cold slicing solution (234 mM sucrose, 11 mM D-glucose, 2.5 mM KCl, 1.25 mM NaH_2_PO_4_, 0.5 mM CaCl_2_, 2.0 mM MgSO_4_, 26 mM NaHCO_3_) (Ma et al., 2015) using forceps. To collect sufficient material for RNA sequencing (~100 ng total RNA) hypothalami were pooled. For each of the four groups (3 and 7 dpf wildtype and mutant embryos) three independent replicates were collected adding up to a total of 12 samples. The collected tissue was preserved in RNA*later* RNA stabilisation solution (Qiagen) until RNA was isolated using RNeasy Mini Kit (Qiagen) according to manufacturer’s instructions. RNA Library preparation was performed by the Core Unit Systems Medicine of the University of Würzburg according to the Illumina TruSeq stranded mRNA Sample Preparation Guide with 100 ng of input RNA and 15 PCR cycles. All 12 libraries were pooled and sequenced on a NextSeq 500 with a read length of 150 nt. Sequenced reads were mapped with the RNA-Seq aligner software STAR (Dobin et al., 2013) to the Ensembl *Danio rerio* genome version GRCz10. Expected read counts were calculated by RSEM (Li and Dewey, 2011). For detection of differentially expressed genes the Bioconductor/R package DESeq2 was used (Love et al., 2014). From our RNA sequencing data we decided to focus on a total of 82 genes relevant for Fgf-signalling, including *fgf* genes, *fgf* receptor genes, Fgf-signalling pathway genes and Fgf downstream target genes. Genes were chosen according to Itoh (2007), Ornitz and Itoh (2015), zfin.org and the KEGG pathway mapping tool (Kanehisa and Goto, 2000). Heat maps with dendrograms were generated using R (package ‘made4’). Genes with a base mean of ≥10 in all four groups were considered as expressed in the hypothalamus. Genes with a base mean ≥10 and a fold change ≥1.5 were considered as differentially expressed. When a gene passed the selection criteria in at least one of the three comparisons (wildtype at 3 vs 7 dpf, wildtype vs. mutant at 3 and 7 dpf) it was represented in the heat map.

### 3D structural analysis

For 3D modelling of the zebrafish Fgf3 structure the amino acid sequence was sent to the Phyre2 server (Kelley et al., 2015). 155 residues from the entire sequence were modelled with a 100% confidence level using pdb entry 1ihk (Plotnikov et al., 2001) and were subsequently used for the structural assessment of the Fgf3 variants. The structure of Fgf3 wildtype protein as well as truncated Fgf3 mutant (*fgf3*^*t24152*^) and morphant protein were modelled bound to a human FGF1 receptor based on the pdb entry 3ojv (Beenken et al., 2012). The structures were visualised with PyMOL (PyMOL Molecular Graphics System, version 2.0, Schrödinger, LLC) and superpositions were performed in coot (Emsley et al., 2010).

### Image acquisition, cell quantification and area measurements

Prior to image acquisition live embryos were mounted in 3% methylcellulose in Danieau’s solution. Embryos processed for whole mount *in situ* hybridisation and immunohistochemistry were mounted in 80% glycerol in PBT. Images were acquired using a stereo microscope (M205FA, Leica) with Leica Application Suit 3.8.0 (Leica) software for live images, a light microscope (Axiophot, Zeiss) with AxioVision Rel. 4.8 software (Zeiss) for whole mount *in situ* hybridisations or a confocal microscope (Eclipse Ti, Nikon) with NIS Elements AR 3.22.15 software (Nikon) for fluorescent stainings.

For cell quantifications and area measurements Fiji 1.8.0 imaging software (Schindelin et al., 2012) with the BioVoxxel Image Processing and Analysis Toolbox (Brocher, 2015) was used. Further, area measurements of the hypothalamic region stained for *nkx2.4b* were obtained through semi-automatic image analysis followed by manual validation and, if required, manual correction of the computed area. All quantifications were done double blinded. Z-stacks of confocal images were collapsed into maximum intensity projections for the figure panels. Background subtraction, as well as brightness and contrast adjustments were performed on the entire image during figure preparation using Fiji 1.8.0 imaging software (Schindelin et al., 2012) with the BioVoxxel Image Processing and Analysis Toolbox (Brocher, 2015) and GIMP 2.10 (The GIMP Team).

### Statistical analysis

Rstudio 1.0.143 (Rstudio, Inc.) was used for statistical analysis and graph design. Median and median absolute deviation (MAD) were calculated for each group and are presented in Suppl. Table 3. Normal distribution was calculated using the Shapiro-Wilk test. If the null hypothesis for normal distribution was accepted a parametric test was used, either a two-sample t-test or a one-way ANOVA depending on number of groups. If the null hypothesis for normal distribution was rejected a non-parametric test was used, either a Mann-Whitney-U test or a Kruskal-Wallis test depending on number of groups. For multiple group comparisons Tukey’s Honest Significance Differences or Nemenyi tests were used as post hoc tests following a one-way ANOVA or a Kruskal-Wallis test, respectively. Results were considered significant with a p-value of 0.05 or lower. Significance levels of 0.05, 0.01 and 0.001 are indicated in the graphs with one, two or three asterisks, respectively.

## Results

### *fgf3* is expressed in the developing hypothalamus and spatially correlates with the location of putative serotonergic CSF-c cells

Among the described Fgf ligands, *fgf3* exhibits the most prominent distribution in the posterior hypothalamus (Bosco et al., 2013; Herzog et al., 2004; Jackman et al., 2004; Liu et al., 2013; Reifers et al., 1998; Topp et al., 2008) presumably where serotonergic precursor cells are located, therefore rendering Fgf3 a likely candidate regulating the development of hypothalamic serotonergic CSF-c cells. To explore the dynamics of hypothalamic *fgf3* expression we performed a spatio-temporal expression analysis. We could confirm presence of *fgf3* transcripts in the developing hypothalamus (Fig. 1). More specifically, transcripts were first detectable at 20 somites in the hypothalamic primordium (Fig. 1. C, D). This expression was maintained until 30 hpf (Fig. 1. E, F). From 36 hpf and onwards the signal was restricted to the posterior hypothalamus, and gradually got limited to cells located medially along the posterior recess of the third ventricle at 68 hpf (Fig. 1G-J). Our expression analysis, thus, strengthens the hypothesised role of Fgf3 as an important regulator during the ontogeny of hypothalamic serotonergic cells. Notably, *fgf3* expression precedes the expression of *tph1a* and 5-HT by several hours (Bellipanni et al., 2002; Bosco et al., 2013) arguing for an early effect on progenitors and possibly differentiation.

### Fgf3 regulates *etv5b* expression in the posterior hypothalamus

The transcription factor Etv5b is a downstream target of Fgf-signalling in various contexts (Mason, 2007; Ornitz and Itoh, 2015; Raible and Brand, 2001; Roehl and Nüsslein-Volhard, 2001; Roussigné and Blader, 2006). However, the identity of the Fgf(s) regulating Etv5b and thereby the ontogeny of hypothalamic serotonergic populations remains unknown. Preliminary observations from Bosco et al. (2013) and our current *fgf3* expression analysis (see above), suggested that Fgf3 might be the main Fgf-ligand in this context. To explore this hypothesis, we investigated hypothalamic *etv5b* expression in the *fgf3* mutant, *fgf3*^*t24152*^, which was suggested to be a null mutant (amorph) (Herzog et al., 2004), as well as in *fgf3* morphants. Our analysis revealed a reduced expression of *etv5b* in mutants compared to wildtype siblings at 36 hpf (Fig. 2A-C). Further, *etv5b* expression was reduced in a dose dependent manner as the *in situ* hybridisation signal was weaker in homozygotes than in heterozygotes. Our RNA sequencing data supported these observations by revealing fewer *etv5b* transcripts in homozygous *fgf3*^*t24152*^ mutants than in wildtypes at 3 and 7 dpf (Fig. 2D), although the read count values did not pass our defined thresholds. Similarly we observed a reduction of *etv5b* expression in *fgf3* morphants compared to controls (Fig. 2E, F). However, the *etv5b* signal was never completely abolished, neither in our *in situ* hybridisation nor in our RNA sequencing experiments. Thus, we showed that Fgf3 regulates *etv5b* expression in the developing posterior hypothalamus.

### Fgf3 impacts on monoaminergic cell development in posterior hypothalamus

After identifying Fgf3 as a possible regulator of hypothalamic serotonergic CSF-c cell development based on expression data, we next tested this hypothesis functionally. For this we applied three complementary strategies to manipulate *fgf3* activity. Firstly, we used the *fgf3*^*t24152*^ mutant, which has a G to A transition point mutation resulting in a premature stop codon and, thereby, a truncated Fgf3 protein with 69.1% of the wildtype amino acid sequence remaining (Suppl. Fig. 1D) (Herzog et al., 2004). Secondly, we created a *fgf3* morpholino knock-down, which leads to a truncated protein containing 49.6% of the wildtype amino acid sequence (Suppl. Fig. 1D). Finally, we applied CRISPR/Cas9 to generate indel mutations causing either a nonsense amino acid sequence or a premature stop close to the N-terminal (Suppl. Fig. 2). All embryos with a manipulated *fgf3*, irrespective of the used strategy, showed similar defects in ear and craniofacial development at 72 hpf (Suppl. Fig. 3). Embryos exhibited fused otoliths presumably due to a role of Fgf3 in anterior ear specification, and malformations of the ventral head skeleton attributed to the loss of ceratobranchial cartilage (Hammond and Whitfield, 2011; Herzog et al., 2004). Additionally, embryos with manipulated *fgf3* displayed a defect in swim bladder development or inflation (not shown). The fact that we could confirm these phenotypes in homozygous *fgf3*^*t24152*^ mutants and reproduce similar defects in *fgf3* morpholino and CRISPR/Cas9 injected embryos show that all three approaches to manipulate *fgf3* produce qualitatively comparable results. Apart from the described defects, no severe morphological abnormalities or increased cell death were observed (Suppl. Fig. 3, 4). Next we focused on the impact on hypothalamic serotonergic CSF-c cell development applying all three strategies in parallel. The number of hypothalamic 5-HT immunoreactive cells in the intermediate (i.) and posterior (p.) populations were quantified at 72 hpf, a stage when the maturation of the serotonergic cells is well on the way (Bellipanni et al., 2002; Bosco et al., 2013; McLean and Fetcho, 2004). In addition, the same embryos were co-stained with a TH1 antibody labelling catecholaminergic cells. TH1-positive cells of the posterior tuberculum/hypothalamus are subdivided into several subpopulations (Rink and Wullimann, 2002). We quantified the TH1 immunoreactive CSF-c cells in regions DC 4/5/6 and DC 7, which are dopaminergic (Yamamoto et al., 2010, 2011). Region DC 4/5/6 is located close to the i. serotonergic population while DC 7 cells are intermingled with the p. serotonergic population (Kaslin and Panula, 2001; McLean and Fetcho, 2004).

**Fig. 3.**
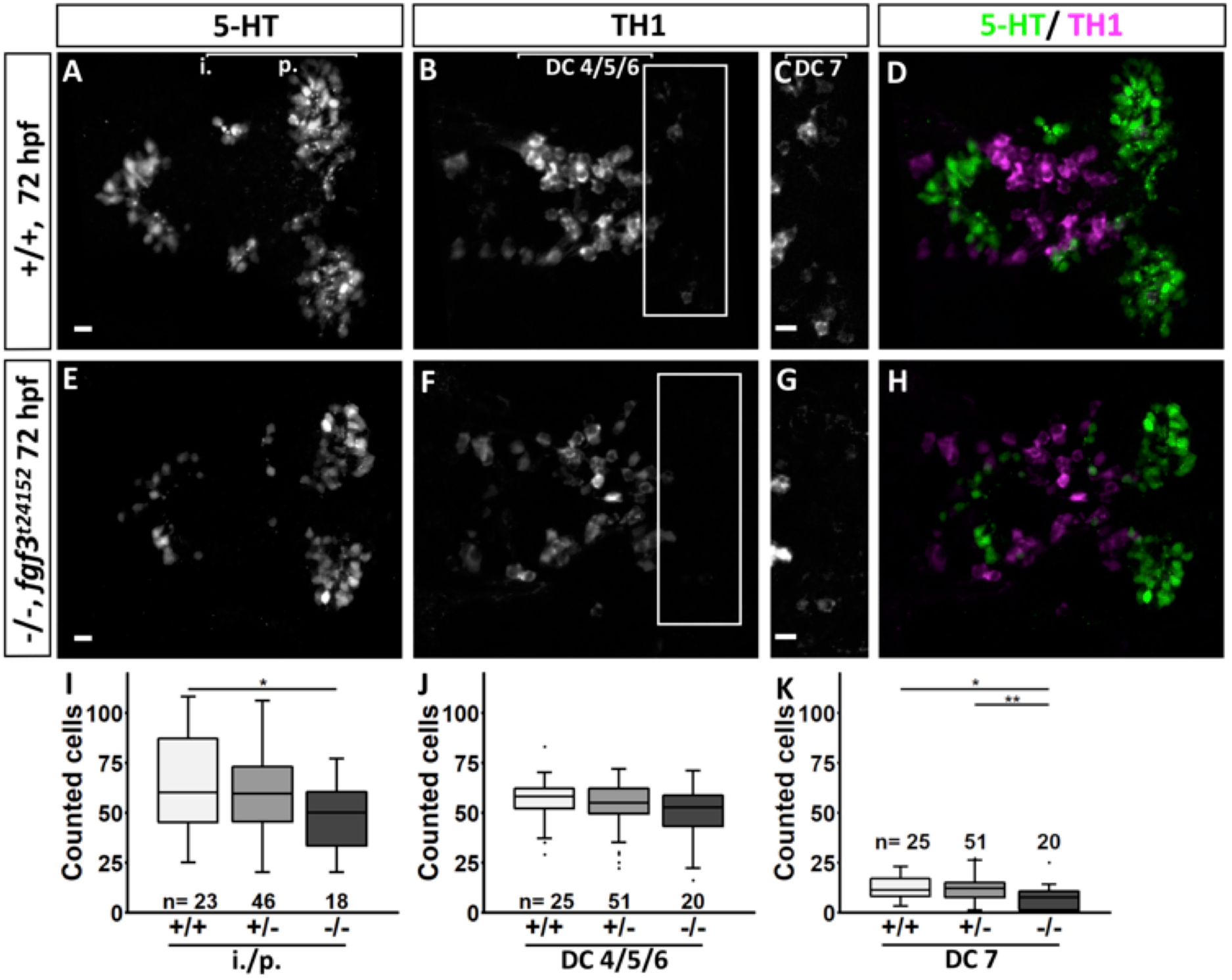
Quantification of the number of serotonergic cells in the intermediate (i.)/posterior (p.) clusters and of dopaminergic cells in the DC 4/5/6 and DC 7 clusters in the hypothalamus of *fgf3*^*t24152*^ mutants at 72 hpf. **(A-H)** Confocal maximum intensity projections from wildtype (+/+) and homozygous *fgf3*^*t24152*^ mutant (-/-) siblings immuno stained for 5-HT (green) and TH1 (magenta) shown as single channels (**A-C, E-G**) and merged (**D, H**). **C** and **G** show boxed areas in **B** and **F**, respectively, with adjusted brightness and contrast to reveal the faint TH1 immunoreactive cells of the DC 7 cluster. Ventral views, anterior to the left. Scale bars = 10 µm. **(I-K)** Quantifications of 5-HT and TH1 positive cells in all three genotypes. The number of serotonergic cells was counted in the i./p. clusters as indicated by the line in **A**. The number of dopaminergic cells was counted in the DC 4/5/6 and DC 7 clusters as indicated by the lines in **B** and **C**, respectively. Boxplots show median, 25-75% percentile, min/max whiskers and outliers depicted as dots. n = number of analysed individuals.

**Fig. 4.**
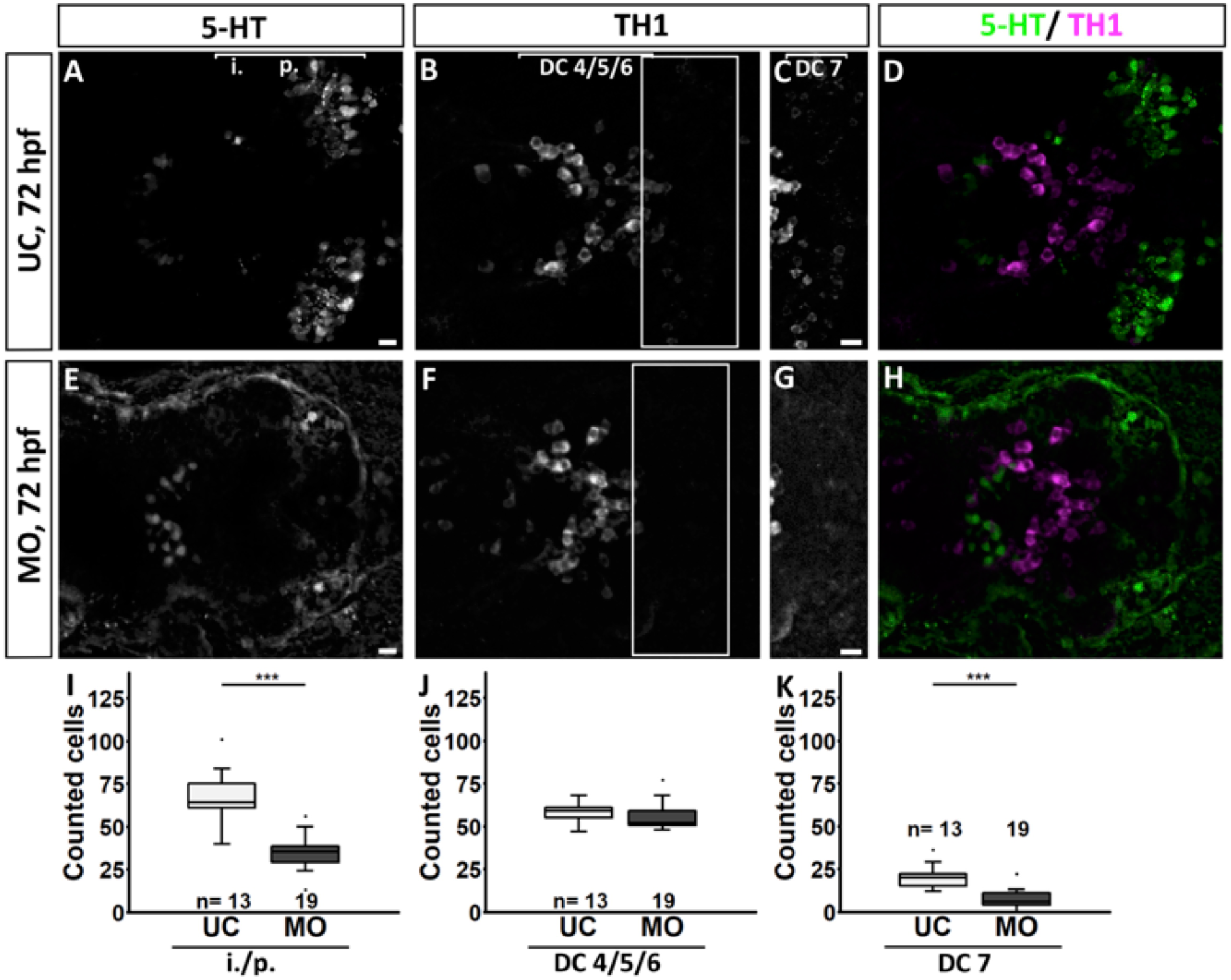
Quantification of the number of serotonergic cells in the intermediate (i.)/posterior (p.) clusters and of dopaminergic cells in the DC 4/5/6 and DC 7 clusters in the hypothalamus of *fgf3* morphants at 72 hpf. **(A-H)** Confocal maximum intensity projections from uninjected control (UC) and morpholino injected (MO) siblings immuno stained for 5-HT (green) and TH1 (magenta) shown as single channels (**A-C, E-G**) and merged (**D, H**). **C** and **G** show boxed areas in **B** and **F**, respectively, with adjusted brightness and contrast to reveal the faint TH1 immunoreactive cells of the DC 7 cluster. Ventral views, anterior to the left. Scale bars = 10 µm. **(I-K)** Quantifications of 5-HT and TH1 positive cells in control and morphant siblings. The number of serotonergic cells was counted in the i./p. clusters as indicated by the line in **A**. The number of dopaminergic cells was counted in the DC 4/5/6 and DC 7 clusters as indicated by the lines in **B** and **C**, respectively. Boxplots show median, 25-75% percentile, min/max whiskers and outliers depicted as dots. n = number of analysed individuals.

We found that homozygous *fgf3*^*t24152*^ mutants, *fgf3* morpholino and CRISPR/Cas9 injected embryos had reduced numbers of serotonergic CSF-c cells, but depending on the used approach the severity of the phenotype varied. Specifically, homozygous *fgf3*^*t24152*^ mutants had 17% fewer serotonergic CSF-c cells than wildtype siblings (Fig. 3, Suppl. Table 3), *fgf3* morphants had a reduction of 45% compared to controls (Fig. 4, Suppl.Table 3), and CRISPR/Cas9 injected embryos had 49% fewer serotonergic CSF-c cells than uninjected controls and 42% fewer than control siblings injected with Cas9 only (Fig. 5, Suppl. Table 3). The reduction of i./p. serotonergic CSF-c cells persisted at 4 dpf in *fgf3* morphants with a loss of 33% (Suppl. Fig. 5, Suppl. Table 3).

**Fig. 5.**
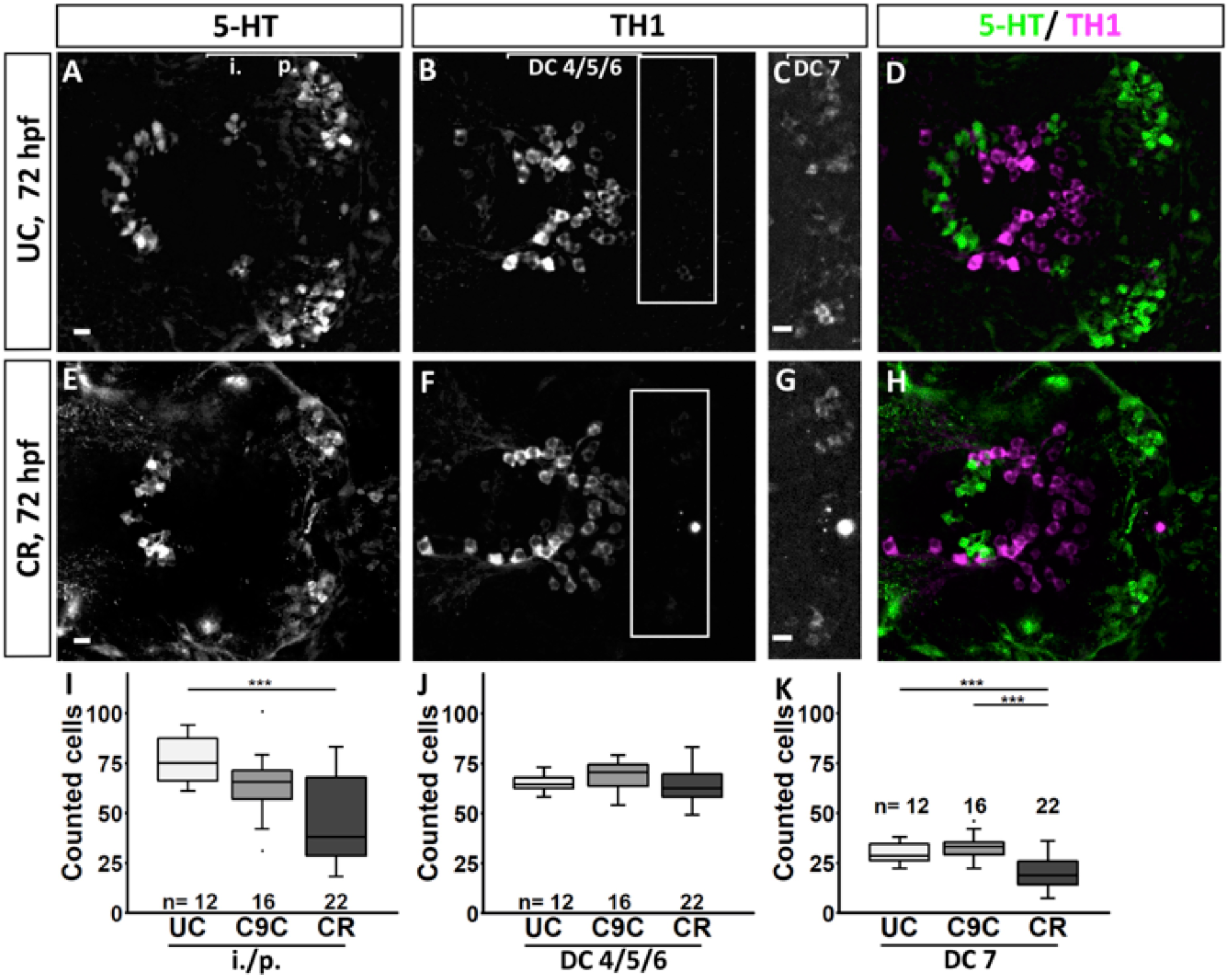
Quantification of the number of serotonergic cells in the intermediate (i.)/posterior (p.) clusters and of dopaminergic cells in the DC 4/5/6 and DC 7 clusters in the hypothalamus of *fgf3* CRISPR/Cas9 injected embryos at 72 hpf. **(A-H)** Confocal maximum intensity projections from uninjected control (UC) and CRISPR/Cas9 injected (CR) siblings immuno stained for 5-HT (green) and TH1 (magenta) shown as single channels (**A-C, E-G**) and merged (**D, H**). **C** and **G** show boxed areas in **B** and **F**, respectively, with adjusted brightness and contrast to reveal the faint TH1 immunoreactive cells of the DC 7 cluster. Ventral views, anterior to the left. Scale bars = 10 µm. **(I-K)** Quantifications of 5-HT and TH1 positive cells in UC, Cas9 only injected (C9C) and CR siblings. The number of serotonergic cells was counted in the i./p. clusters as indicated by the line in **A**. The number of dopaminergic cells was counted in the DC 4/5/6 and DC 7 clusters as indicated by the lines in **B** and **C**, respectively. Boxplots show median, 25-75% percentile, min/max whiskers and outliers depicted as dots. n = number of analysed individuals.

With respect to the TH1 immunoreactive cells, the numbers in DC 4/5/6 were never significantly affected after any of the *fgf3* manipulations (Fig. 3J, 4J, 5J, Suppl. Fig. 5J, Suppl. Table 3). In contrast, homozygous *fgf3*^*t24152*^ mutants, *fgf3* morpholino and CRISPR/Cas9 injected embryos had fewer cells in DC 7. A reduction of TH1-positive cells by 32% was detectable in homozygous *fgf3*^*t24152*^ mutants compared to wildtype siblings (Fig. 3, Suppl. Table 3). Further, homozygous *fgf3*^*t24152*^ mutants had 38% fewer TH1-positive cells than heterozygotes. In *fgf3* morphants, the number of TH1-expressing cells in DC7 was reduced by 70% compared to uninjected controls (Fig. 4, Suppl. Table 3). CRISPR/Cas9 injected embryos had 35% fewer TH1-positive cells in comparison to uninjected controls, and 44% fewer compared to controls injected with Cas9 only (Fig. 5, Suppl. Table 3). The reduction of TH1-positive DC 7 cells persisted at 4 dpf in *fgf3* MO embryos with a loss of 34% (Suppl. Fig. 5, Suppl. Table 3).

Taken together, using three independent techniques to manipulate *fgf3* activity, we demonstrated a consistent loss of monoaminergic CSF-c cells after *fgf3* impairment, showing a developmental dependency of these cells on Fgf3. Interestingly, this seemed to specifically affect the populations in the posterior hypothalamus.

### oxytocin (*oxt)* and *cortistatin* (*cort)* expressing neuroendocrine cells are unaffected by impaired *fgf3* function

To test the specific dependence of the monoaminergic CSF-c cells in the posterior hypothalamus on Fgf3, three additional cell populations, including neuroendocrine cells expressing *oxytocin (oxt), arginine vasopressin (avp)* and *cortistatin (cort)* were investigated (Devo set al., 2002; Eaton et al., 2008; Unger and Glasgow, 2003). These populations were chosen due to their spatial proximity to the hypothalamic monoaminergic regions. *fgf3*^*t24152*^ mutants and *fgf3* morphants were labelled for *oxt, avp* and *cort* at 72 hpf, and the number of positive cells was counted. The *oxt* and *cort* cell numbers were not decreased in hetero- and homozygous *fgf3*^*t24152*^ mutants (Fig. 6, Suppl. Table 3) or *fgf3* morphants (Fig. 7, Suppl. Table 3) compared to control siblings. However, there was a significant reduction in the number of *avp*-positive cells. Homozygous *fgf3*^*t24152*^ mutants lost 16% of the cells compared to wildtypes and 14% compared to heterozygotes (Fig. 6H, Suppl. Table 3). In *fgf3* morphants, *avp*-expressing cell numbers decreased by 11% compared to controls (Fig. 7H, Suppl. Table 3). Thus, *avp*-expressing, but not *oxt-* or *cort*-expressing cells showed dependency on Fgf3. Interestingly, the most posteriorly located *avp*-positive cells were more affected than anteriorly located cells in mutants and morphants (Fig. 6E, 7E), and therefore mirroring the effects on the TH1-positive cells.

**Fig. 6.**
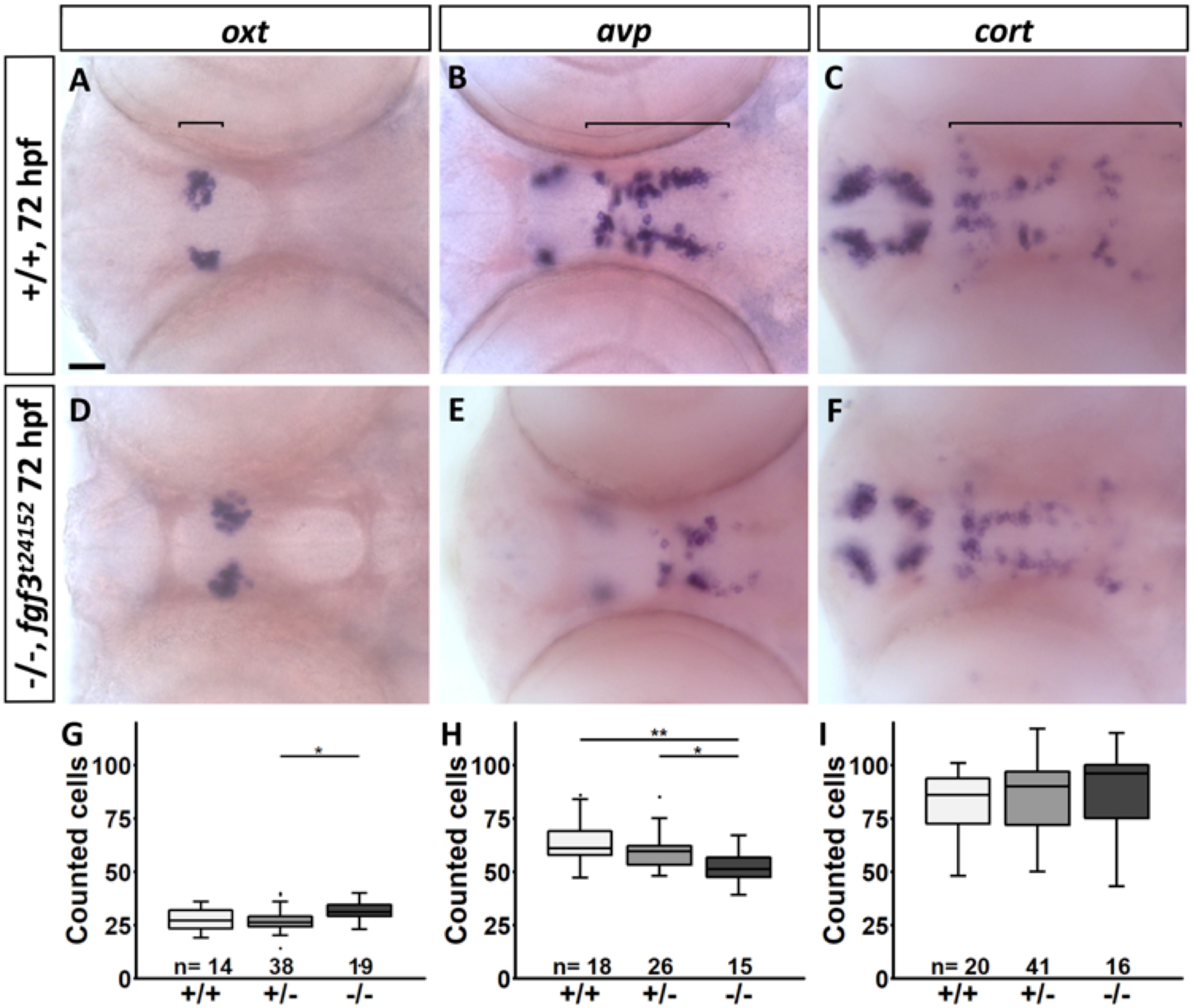
Quantification of the number of *oxt, avp* and *cort* expressing cells in the hypothalamus of *fgf3*^*t24152*^ mutants at 72 hpf. **(A-F)** Light microscopic pictures of wildtype (+/+) and homozygous *fgf3*^*t24152*^ mutant (-/-) siblings processed for RNA *in situ* hybridisation. Note that the posteriorly located *avp* expressing cells are more affected by impaired *fgf3* than the anterior ones. Ventral views, anterior to the left. Scale bars = 30 µm. **(G-I)** Quantifications of *oxt-, avp-* and *cort*-positive cells in all three genotypes. Cell clusters used for the analyses are indicated by the lines in **A-C**. Boxplots show median, 25-75% percentile, min/max whiskers and outliers depicted as dots. n = number of analysed individuals.

**Fig. 7.**
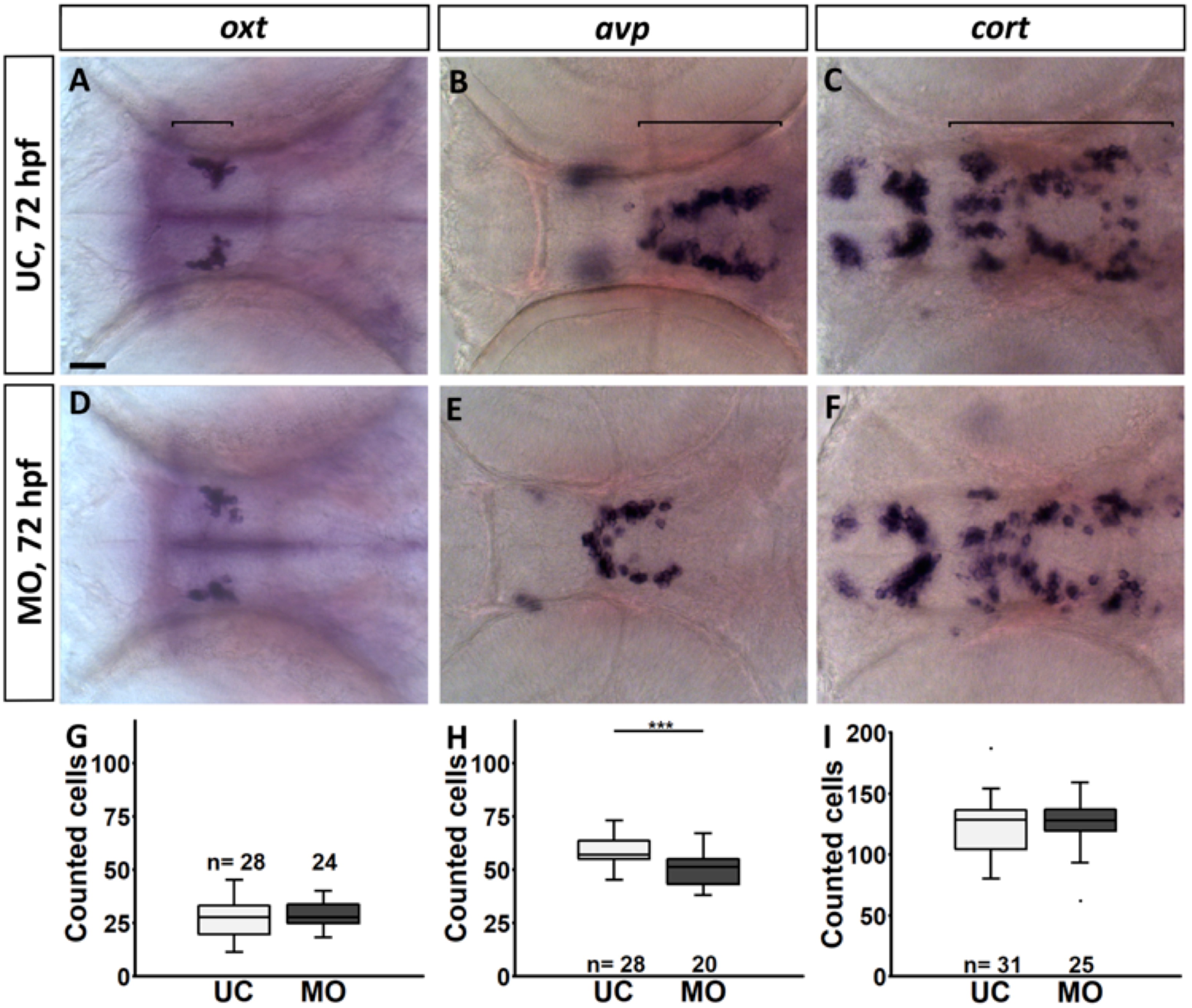
Quantification of the number of *oxt, avp* and *cort* expressing cells in the hypothalamus of *fgf3* morphants at 72 hpf. **(A-F)** Light microscopic pictures of uninjected control (UC) and morpholino injected (MO) siblings processed for RNA *in situ* hybridisation. Note that the posteriorly located *avp* expressing cells are more affected by impaired *fgf3* than the anterior ones. Ventral views, anterior to the left. Scale bars = 30 µm. **(G-I)** Quantifications of *oxt-, avp-* and *cort*-positive cells in control and morphant siblings. Cell clusters used for the analyses are indicated by the lines in **A-C**. Boxplots show median, 25-75% percentile, min/max whiskers and outliers depicted as dots. n = number of analysed individuals.

### *fgf3*^*t24152*^ mutants and *fgf3* morphants have a smaller hypothalamus

To determine whether the loss of hypothalamic monoaminergic CSF-c cells, as well as *avp*-expressing cells consequently resulted in a smaller hypothalamus, we measured the size of the hypothalamic domain in *fgf3^t24152^* mutants and *fgf3* morphants. *nkx2.4b* expression at 36, 48 and 72 hpf was used to visualise the hypothalamic domain, which was subsequently measured in a semi-automatic manner. In homozygous *fgf3*^*t24152*^ mutants a trend (6%) towards a smaller hypothalamic domain was observed at 36 hpf (Fig. 8, Suppl. Table 3). At 48 hpf a significant reduction of 11% in homozygous *fgf3*^*t24152*^ mutants compared to wildtype siblings was recorded, and at 72 hpf the hypothalamus was smaller in homozygous *fgf3*^*t24152*^ mutants compared to both heterozygous mutant (10%) and wildtype (14%) siblings. As expected, measurements of *fgf3* morphants revealed significantly reduced hypothalami. At 36 hpf the *nkx2.4b*-positive domain in *fgf3* morphants was 19% smaller compared to uninjected controls (Fig. 9, Suppl. Table 3). Further, at 48 and 72 hpf the reduction was 16% and 15%, respectively. Thus, we noticed that the size of the hypothalamic domain was more reduced in *fgf3* morphants than in *fgf3*^*t24152*^ mutants at all stages examined, which was in line with the stronger monoaminergic phenotypes observed in *fgf3* morphants compared to *fgf3*^*t24152*^ mutants described above.

**Fig. 8.**
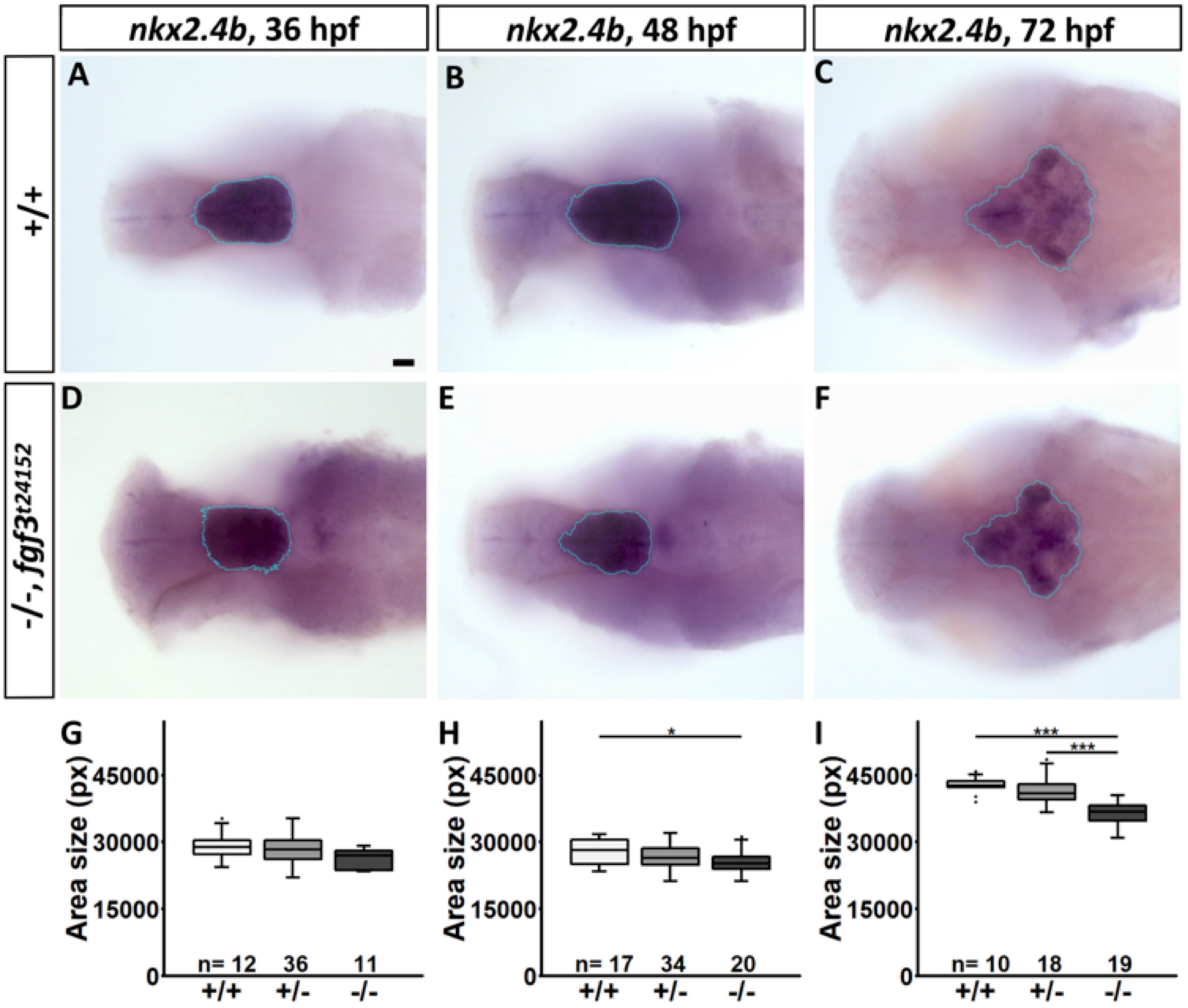
The size of the hypothalamic *nkx2.4b* domain is significantly reduced in *fgf3*^*t24152*^ mutants. **(A-F)** Light microscopic pictures showing expression of *nkx2.4b* in wildtype (+/+) and homozygous *fgf3*^*t24152*^ mutant (-/-) siblings at 36, 48 and 72 hpf visualised by RNA *in situ* hybridisation. Outline of semi-automated measurement of hypothalamic area is highlighted in blue. Ventral views, anterior to the left. Scale bar = 30 µm. **(G-I)** Area measurements (pixels) in embryos with either of the three genotypes. Boxplots show median, 25-75% percentile, min/max whiskers and outliers depicted as dots. n= number of analysed individuals.

**Fig. 9.**
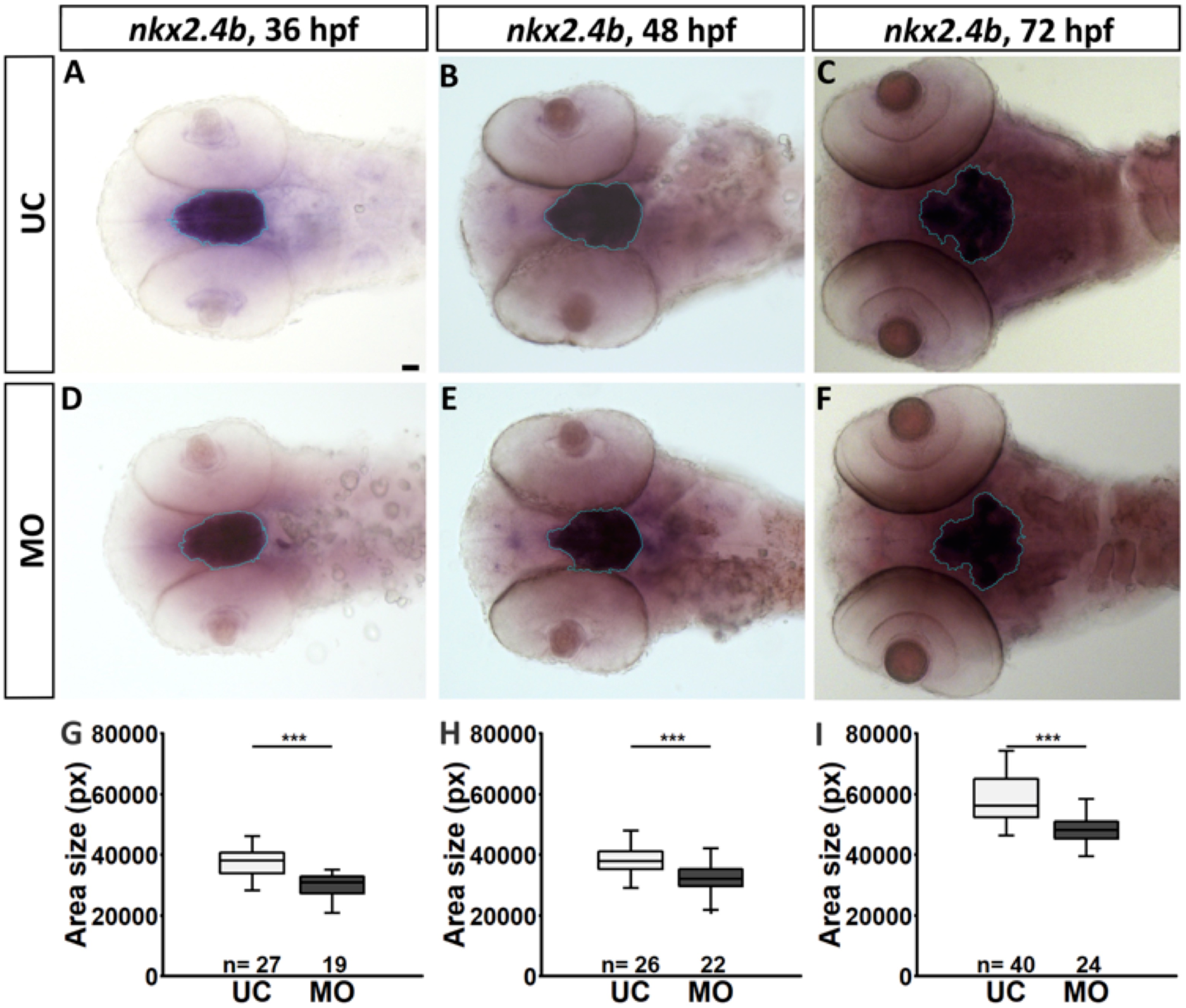
The size of the hypothalamic *nkx2.4b* domain is significantly reduced in *fgf3* morphants. **(A-F)** Light microscopic pictures showing expression of *nkx2.4b* in uninjected control (UC) and morpholino injected (MO) siblings at 36, 48 and 72 hpf visualised by whole mount RNA in situ hybridisation. Outline of semi-automated measurement of hypothalamic area is highlighted in blue. Ventral views, anterior to the left. Scale bar = 30 µm. **(G-I)** Area measurements (pixels) in control and morphant siblings. Boxplots show median, 25-75% percentile, min/max whiskers and outliers depicted as dots. n= number of analysed individuals.

### *fgf3*^*t24152*^ mutant and morphant Fgf3 may still interact with Fgf receptors

To estimate the impact of the *fgf3*^*t24152*^ mutation and the exon2/intron2 splice blocking morpholino on the folding, stability and receptor binding capacity of the truncated isoforms of Fgf3 we carried out computational 3D modelling comparing both truncated isoforms to wildtype Fgf3 (Fig. 10). As described above, the *fgf3*^*t24152*^ mutation resulted in a Fgf3 isoform with about 70% of the wildtype amino acid sequence remaining, while the splice morpholino generated an isoform with about 50% of the wildtype sequence intact (Suppl. Fig. 1D). Notably, receptor binding is likely to still be possible for the *fgf3*^*t24152*^ mutant Fgf3 since a major portion of the interfaces with the Fgf receptor may remain intact (Fig. 10B). As expected the 3D model of the morpholino knock-down isoform of Fgf3 showed less preserved structure than the *fgf3*^*t24152*^ isoform and therefore a lower probability to interact with the receptor (Fig. 10C). Both isoforms are likely to be less stable than wildtype Fgf3.

**Fig. 10.**
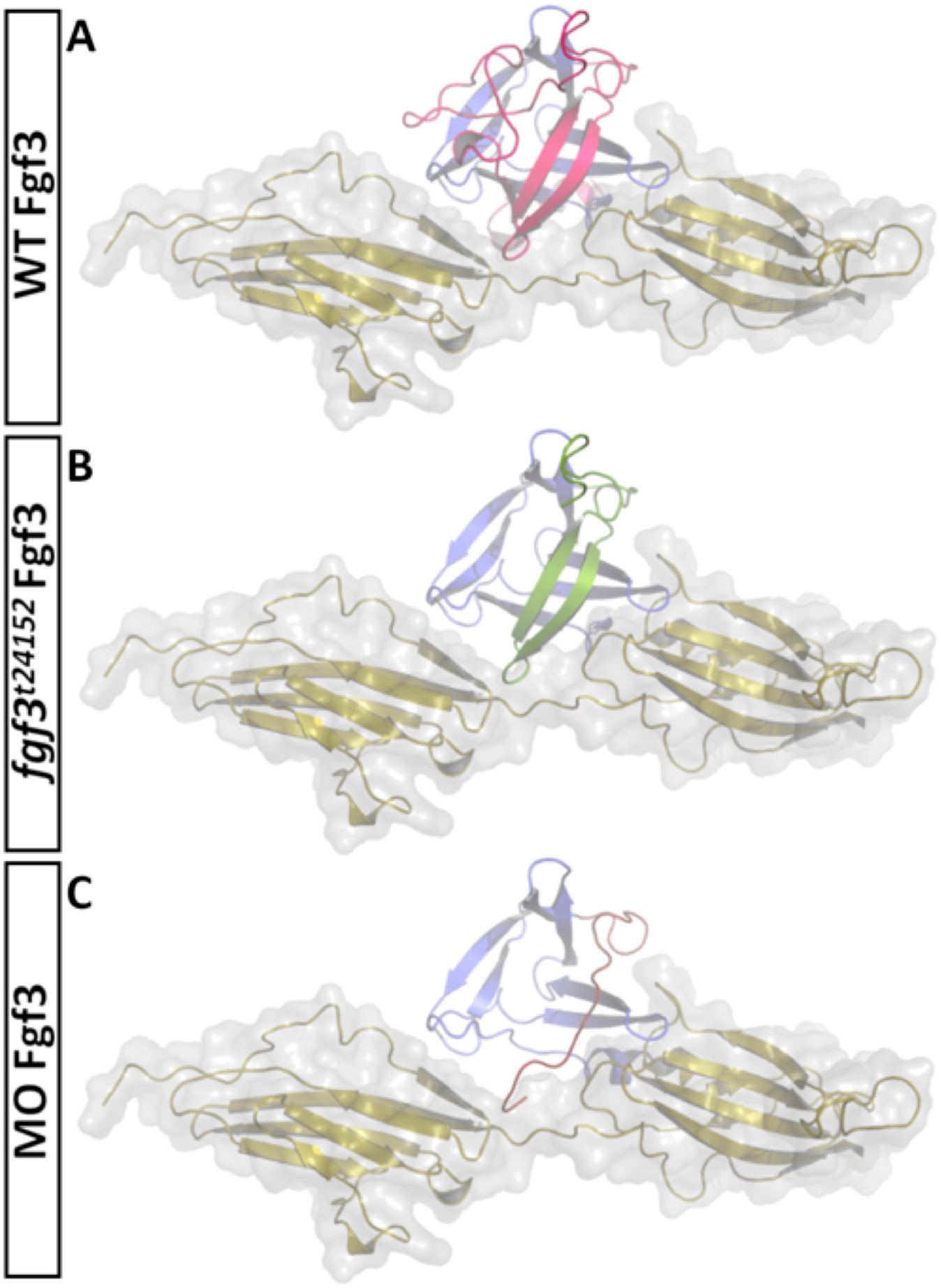
Ribbon representation of models of zebrafish Fgf3 isoforms bound to a FGF1 receptor. **(A)** Ribbon representation of a zebrafish Fgf3 model (blue and pink) bound to a FGF1 receptor (yellow). The overall part affected by the *fgf3*^*t24152*^ mutation or the *fgf3* morpholino is colored in pink. **(B)** The deleted region of the *fgf3*^*t24152*^ mutant isoform is omitted. The part that remains (green) as compared to the morpholino knock-down isoform still folds. Notably receptor binding would be still possible if the variant maintains partial folding, since most of the interfaces with the FGF1 receptor would stay intact. **(C)** Model of the morpholino knock-down model. The exchanged sequence is colored in red. In comparison to the *fgf3*^*t24152*^ mutant model it lacks even more structural elements rendering a correct folding and residual receptor interactions less likely.

**Fig. 11.**
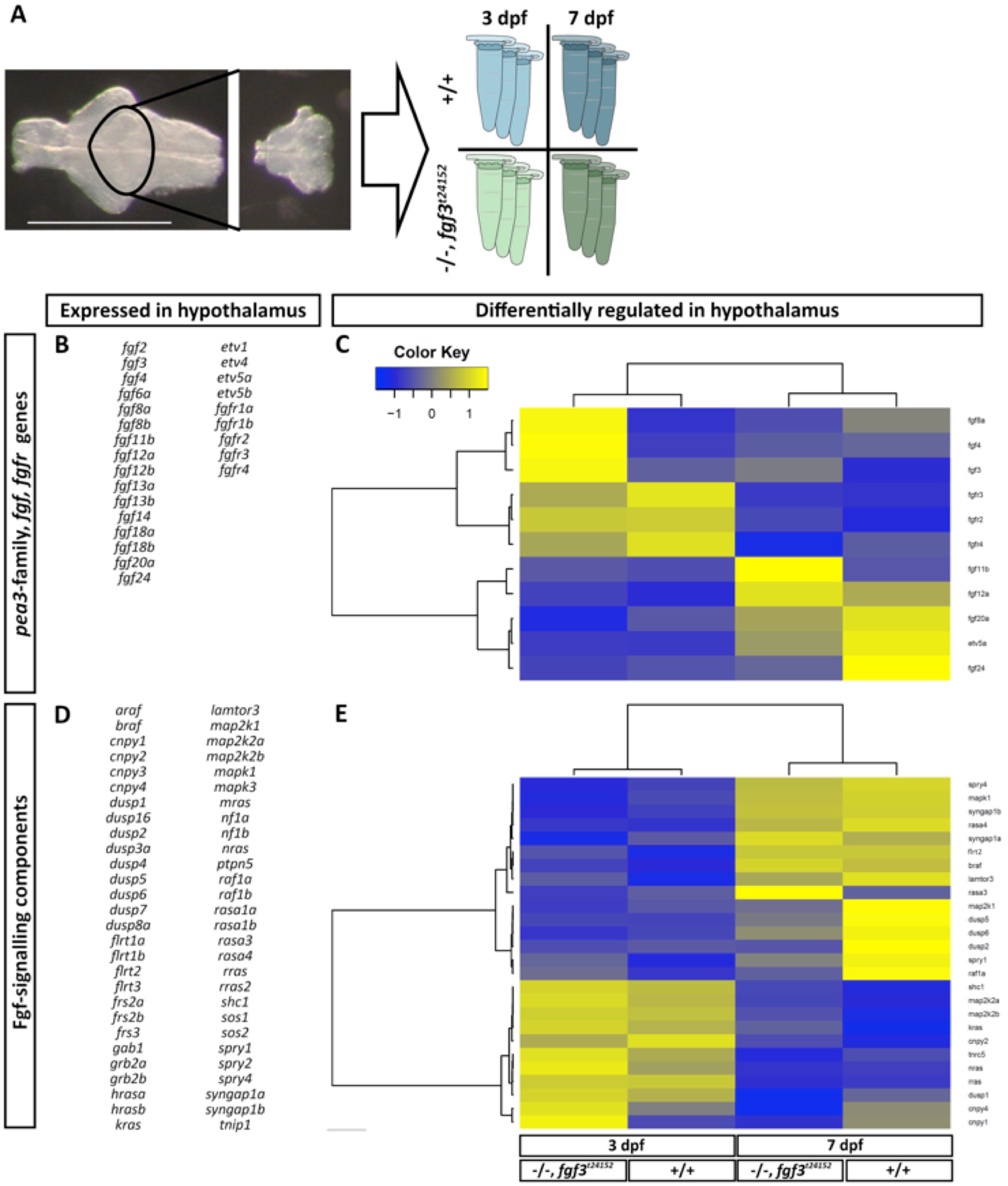
Hierarchical clustering of hypothalamic Fgf-signalling pathway genes and downstream targets analysed in *fgf3*^*t24152*^ mutants (-/-) and wildtypes (+/+) at 3 and 7 dpf. **(A)** Scheme illustrating sample preparation and groups used for RNA sequencing. Micrographs show the dissected hypothalamic area (ventral view, anterior to the left) collected from wildtypes (blue tubes) and *fgf3*^*t24152*^ mutants (green tubes) at 3 (light tubes) and 7 dpf (dark tubes). For each group three independent tissue collections were performed resulting in a total of 12 samples processed for RNA sequencing. Scale bar = 500 µm **(B, D)** List of *fgfs, fgfrs*, Fgf downstream target genes and Fgf-signalling components expressed (base mean ≥10) in the hypothalamus of -/- and +/+ embryos. All genes except *shc2* pass the base mean threshold in all four groups. *shc2* does not pass the base mean threshold in the 3 dpf -/- group. **(C, E)** Heat maps of differentially regulated (base mean ≥10, fold change ≥1.5) *fgfs*, *fgfrs* and Fgf downstream target genes (**C**) as well as Fgf-signalling components (**E**) in the hypothalamus of -/- and +/+ embryos. Columns represent z-score of mean values of replicates for each analysed group and rows depict individual genes. Colour key displays z-score ranging from ≥ −1.5 to ≤ 1.5. Colour intensity represents expression levels of a gene for each group with blue or yellow indicating low or high expression, respectively.

### *fgf3^t24152^* mutants exhibit minor alterations in the hypothalamic fgf transcriptome

The paracrine family of Fgf ligands include multiple members in addition to Fgf3 (Itoh, 2007). Further, Fgf-signalling contains numerous regulatory feedback systems (Ornitz and Itoh, 2015). Therefore, it may be that one or several previously overseen Fgfs and/or that the feedback systems compensate for a loss of Fgf3 functionality. Such compensatory mechanisms could in turn explain why some of the posterior monoaminergic CSF-c and *avp*-expressing cells remain after *fgf3* impairment. To test this possibility we accomplished a transcriptome analysis.

RNA sequencing was performed on dissected hypothalami of homozygous *fgf3*^*t24152*^ mutants and wildtype cousins at 3 and 7 dpf (Fig. 11A). Our data analyses of 31 known zebrafish *fgf* genes (Suppl. Table 4) revealed that transcripts for other previously overseen *fgfs* are present in wildtypes and mutants (Fig. 11B). In addition to *fgf3, fgf4* and *fgf8a* transcripts, which were already found in the developing hypothalamus (Herzog et al., 2004; Jackman et al., 2004; Liu et al., 2013; Reifers et al., 1998), we detected 14 further *fgf* transcripts in all groups, including *fgf2, fgf4, fgf6a, fgf8b, fgf11b, fgf12a, fgf12b, fgf13a, fgf13b, fgf14, fgf18a, fgf18b, fgf20a, fgf24*. Of these *fgf3, fgf12a, fgf20a* and *fgf24* were differentially expressed at 3 and 7 dpf in the wildtype (Fig. 11C). Interestingly, of the 16 detected *fgfs* only *fgf3* was upregulated in mutants at both 3 and 7 dpf compared to wildtypes of the same age (Fig. 11C), which indicates a self-compensatory mechanism of *fgf3. fgf11b* and *fgf24* were up- and downregulated, respectively, in mutants compared to wildtypes, but only at 7 dpf (Fig. 11C). All other detected *fgfs* did not pass our 1.5 fold change criteria for the mutant versus wildtype comparisons. Moreover, our RNA sequencing data showed that all 5 *fgfr* genes were expressed in the hypothalamus of all groups (Fig. 11B). In the wildtype groups *fgfr2, fgfr3* and *fgfr4* were downregulated at 7 dpf compared to 3 dpf (Fig. 11C). Four out of four selected ETS-domain transcription factors, including *etv1, etv4, etv5a* and *etv5b*, were expressed in the hypothalamus of all groups (Fig. 11B). Of these ETS-domain factors only *etv5a* expression increased beyond our predefined fold change threshold from 3 to 7 dpf in wildtypes, but did not change when comparing mutants to wildtypes at 3 or 7 dpf (Fig.11C). RNA sequencing results of Fgf-signalling pathway genes revealed 56 out of 62 selected genes (Suppl. Table 4) to be expressed in the hypothalamus of wildtypes and mutants at 3 and 7 dpf (Fig. 11D). None of these 56 genes were differentially expressed when comparing mutants and wildtypes at 3 dpf, but at 7 dpf *dusp1, dusp2* and *dusp5* were downregulated in the mutants (Fig. 11E).

In summary, our RNA sequencing data showed that transcripts for previously overseen Fgf ligands are present in the developing wildtype hypothalamus. However, except for *fgf3* and *fgf11b*, neither of these were notably differentially regulated in the *fgf3*^*t24152*^ mutant. Furthermore, the alterations in the downstream Fgf-signalling pathway were moderate. This suggests that only minor compensatory mechanisms are at hand in the *fgf3*^*t24152*^ mutant and/or that the truncated Fgf3 isoform resulting from the *fgf3*^*t24152*^ mutation is still, at least partly, functional.

## Discussion

In this study we identify Fgf3 as the main Fgf ligand regulating the ontogeny of monoaminergic CSF-c and of *avp*-expressing cells in the zebrafish posterior hypothalamus. The requirement of Fgf3 follows a posterior to anterior gradient. Further, we show that Fgf3 controls expression of the ETS-domain transcription factor *etv5b*. Based on our current observations that *fgf3* expression precedes that of *tph1a* and 5-HT by several hours (Bellipanni et al., 2002; Bosco et al., 2013), and that the *nkx2.4b*-positive hypothalamic domain is smaller after *fgf3* impairment before the appearance of mature serotonergic CSF-c cells we propose that Fgf3 is critical during early stages when progenitors are still proliferating. This is in line with our earlier finding that Fgf-signalling, via Etv5b, influences the proliferation of hypothalamic serotonergic CSF-c progenitor cells (Bosco et al., 2013). Furthermore, our present study provides evidence for the expression of previously overseen *fgfs* in the developing hypothalamus. We show that the expression of *fgf3* is upregulated upon impairment of *fgf3*, suggesting activation of a self-compensatory mechanism. Together these findings highlight Fgf-signalling, in particular Fgf3, in a novel context as part of a signalling pathway of critical importance for hypothalamic development. Our results have implications for the understanding of vertebrate hypothalamic evolution.

### Fgf3 is present in the posterior hypothalamus and likely acts as a morphogen regulating the expression of ETS-domain transcription factors

Fgf-signalling, acting via Etv5b, is an important pathway for development of serotonergic CSF-c cells in the posterior hypothalamus (Bosco et al., 2013) as well as in other contexts (Mason, 2007; Ornitz and Itoh, 2015; Raible and Brand, 2001; Roehl and Nüsslein-Volhard, 2001; Roussigné and Blader, 2006). However, the Fgf ligands are many and their interactions with Fgfrs are promiscuous (Ornitz and Itoh, 2015). To identify the Fgf ligand active in the developing posterior hypothalamus we searched existing expression data and found that *fgf3* is present in its most posterior region (Herzog et al., 2004), an area that contains serotonergic CSF-c cells (Lillesaar, 2011; McLean and Fetcho, 2004). Thus, we hypothesised that Fgf3 is the main ligand responsible for Fgf activity in this particular region of the brain. As observed elsewhere (Herzog et al., 2004; Liu et al., 2013), we confirmed that *fgf3* is expressed in the developing hypothalamus. Initially it has a broad distribution in the hypothalamic primordium, The expression then gradually becomes limited to the most posterior end in cells located around the posterior recess. Conforming to Fgf3 activity in the posterior hypothalamus, the Fgf responsive downstream targets *etv4, etv5a* and *etv5b* are present there (Bosco et al., 2013). However, in contrast to *fgf3* their expression is not limited to cells at the ventricle, but shows a broader distribution including cells in the parenchyme, thus, fitting the role of Fgf3 as a morphogen (Bökel and Brand, 2013; Ornitz and Itoh, 2015). To test if Fgf3 regulates *etv5b* expression in the hypothalamus we analysed *etv5b* expression in *fgf3*^*t24152*^ mutants and *fgf3* morphants. We noticed a reduction of *etv5b* transcripts in the posterior hypothalamus. These results demonstrate that Fgf3 is an important Fgf ligand in the developing posterior hypothalamus and that Fgf3 positively regulates *etv5b* expression in this brain region. Based on the distribution of transcripts it can be assumed that Fgf3 is secreted by a limited population of cells located at the ventricle, and reaches responsive cells expressing *etv5b* in more lateral positions.

### Fgf3 activity is critical for monoaminergic CSF-c and *avp*-expressing cells in the posterior hypothalamus

Hypothalamic serotonergic progenitors are proliferating at around 36 hpf, and differentiated serotonergic CSF-c cells are detectable at 62 hpf in the i./p. cluster (Bosco et al., 2013; McLean and Fetcho, 2004). Further, Etv5b regulates the proliferation of serotonergic progenitors, and ulitmately the number of mature serotonergic CSF-c cells (Bosco et al., 2013). These observations, in combination with our current finding that Fgf3 impacts on *etv5b* expression, prompted us to investigate whether impairment of *fgf3* would reduce the number of serotonergic CSF-c cells. By impairing *fgf3* we show that, indeed, serotonergic CSF-c cells depend on Fgf3. Irrespective of used approach, we always observed a reduction of hypothalamic serotonergic CSF-c cells. However, the population was never completely abolished, suggesting that some cells are independent of Fgf3, that our loss-of-function approaches result in only partial *fgf3*/Fgf3 impairment and/or that other Fgfs can partly compensate for the loss of Fgf3 (for further details see Discussion below).

In the hypothalamus, dopaminergic CSF-c cells located in region DC 4/5/6 and DC 7 are situated next to or intermingled with 5-HT immunoreactive cells of the i. and p. populations, respectively (Kaslin and Panula, 2001; McLean and Fetcho, 2004; Rink and Wullimann, 2002). Dopaminergic and serotonergic cells belong to the monoaminergic systems and accordingly share the expression of metabolic pathway genes, such as *ddc, mao* and *slc18a2* (*vmat2)* (Yamamoto and Vernier, 2011). This may suggest a common developmental programme for hypothalamic monoaminergic populations, and therefore a mutual dependence on Fgf3. To test if the dopaminergic DC 4/5/6 and DC 7 cells require Fgf3 we quantified them after *fgf3* impairment. Interestingly, we noticed a significant reduction only in the posteriorly located DC 7 population. Similarly, another study showed fewer dopaminergic cells specifically in the DC 7 population of *fgf3*^*t24152*^ mutants (Koch et al., 2014). In contrast, neither the DC4/5/6 nor the DC 7 populations are affected in *etv5b* morphants (Bosco et al., 2013). To further expand the analysis of hypothalamic cell populations potentially affected by Fgf3 we investigated neuroendocrine cells expressing *oxt, avp* and *cort* (Devos et al., 2002; Eaton et al., 2008; Unger and Glasgow, 2003). Of these neuroendocrine populations, the *oxt-* and *cort*-expressing ones were unaffected in *fgf3*^*t24152*^ and *fgf3* morphants. Earlier quantifications of *oxt-* and *cort*-expressing cells in *etv5b* morphants (Bosco et al., 2013) are similar to our present results after *fgf3* impairment. Hence, we conclude that those two neuroendocrine populations are neither dependent on Etv5b nor Fgf3. In contrast, the number of *avp*-expressing cells was reduced in mutants and morphants suggesting that *avp*-expressing cells require Fgf3. Notably, the posterior *avp*-expressing cells were more affected than the anterior ones, similar to our observations of the dopaminergic DC 7 and DC 4/5/6 cells, thus supporting the hypothesis that Fgf3 plays a role predominantly in the posterior hypothalamus. Further, the *avp*-expressing population appears to be *etv5b* independent (Bosco et al., 2013). To summarise, the monoaminergic as well as *avp*-expressing cell populations require Fgf3. However, when comparing the different populations they exhibit distinct Fgf-signalling profiles; 1) *etv5b* is essential only for the posterior serotonergic cells, and 2) there is a posterior to anterior Fgf3 dependence gradient with the highest requirement seen in the posterior populations. We cannot exclude that *etv5b*-independent cell types use another ETS-domain transcription factor activated by Fgf3. The presence of *etv5a* and *etv4* transcripts in this region (Bosco et al., 2013) renders such a scenario possible.

It has been proposed that species-specific developmental programmes may result in “hypothalamic modules”, which can be gained or lost during evolution(Xie and Dorsky, 2017). One striking example of such a putative module is the posterior paraventricular organ surrounding the posterior recess, and which is housing the monoaminergic CSF-c cells. This particular structure is found in teleosts, but absent in tetrapods (Xavier et al., 2017). Thus, it is plausible that Fgf3 and Etv5b are part of a developmental signalling programme promoting the formation of a posterior hypothalamic module.

### Fgf3 likely regulates proliferation of monoaminergic progenitors

The monoaminergic CSF-c cells constitute a considerably large population of cells in the posterior hypothalamus. We therefore hypothesised that loss of those cells, as seen after *fgf3* impairment, would be associated with a smaller hypothalamus. As expected we observed a reduction of the *nkx2.4b*-positive domain at 72 hpf, a stage when numerous monoaminergic cells are present. Interestingly, the size reduction was observable at stages before the appearance of mature monoaminergic cells. This suggests that Fgf3 activity plays a role prior to complete cell maturation, presumably at a stage when monoaminergic progenitors are still proliferating, which is in line with the proposed function of Etv5b (Bosco et al., 2013). It may be argued that the reduction of hypothalamic size is caused by an increased cell death. However, no morphological signs of increased cell death were observed in the *fgf3* impaired embryos (mutants, morphants or CRISPR/Cas9) at any stage investigated, and no notable increase of acridine orange-positive cells was observed in *fgf3* morphants at 24 hpf speaking against this possibility.

### The *fgf3*^*t24152*^ mutant exhibits a milder phenotype than *fgf3* morphants or *fgf3* CRISPR/Cas9 injected embryos

All three strategies (mutant, morpholino and CRISPR/Cas9) to interrupt *fgf3* function resulted in a reduction of monoaminergic CSF-c cells, with up to about 50% loss of serotonergic CSF-c cells on average in *fgf3* CRISPR/Cas9 injected embryos. However, we never saw a complete loss of these cells. This may be explained by 1) some wildtype Fgf3 remaining after manipulation, 2) that the *fgf3*^*t24152*^ allele is not a amorph, hence, not leading to a complete loss of Fgf3 activity, 3) compensatory mechanisms (upregulation of *fgf3* itself, upregulation of other *fgfs*, alterations in the downstream Fgf-signalling pathway and/or in its feedback regulators) and/or 4) that some serotonergic cells develop independently of Fgf3.

Our 3D models of the proteins resulting from the *fgf3^t24152^* mutation and the morpholino knock-down suggest that both isoforms are less stable than the wildtype protein, with the morphant protein being the least stable. However, according to the 3D models showing the ligand/receptor complex both truncated isoforms may still interact with Fgfrs. Again the morphant form exhibits the more severe alteration. It should be pointed out that in the morphant, wildtype transcripts are detectable, and it is therefore likely that we do not have a complete loss-of-function. The *fgf3*^*t24152*^ allele was published as being a nonsense mutation likely to be amorphic leading to a truncated Fgf3 protein with a complete loss of activity (Herzog et al., 2004). Considering this, we expected the homozygous *fgf3*^*t24152*^ mutant to exhibit a more severe hypothalamic phenotype than we observed. With the 3D models and our phenotypic characterisation of the *fgf3*^*t24152*^ mutant at hand we question the previous conclusion that the *fgf3*^*t24152*^ mutation is amorphic, and propose that the *fgf3*^*t24152*^ allele rather corresponds to a hypomorphic mutation.

Our CRISPR/Cas9 experiments were performed in the F0 generation and the mutations will therefore consist of various indels and have a mosaic distribution. The target for our exon 1 guide RNA is situated 153 base pairs from the start codon. In the most severe indel mutation scenario we therefore expect to lose about 80% of the wildtype amino acid sequence. As can be expected in F0 injected embryos, we saw a more variable strength of the phenotype between individuals compared to *fgf3*^*t24152*^ mutants or morphants. In some individuals we noticed an almost complete loss of the posterior monoaminergic cell populations, which correlated with a strongly reduced posterior hypothalamus (Fig. 5D, H). Although we cannot finally conclude from our current data, it is likely that such individuals have lost most, if not all, functional Fgf3. In future studies of stable *fgf3* CRISPR lines it will be interesting to see if there is a complete loss of the posterior hypothalamic CSF-c cells.

Our RNA sequencing results showed that several *fgf* genes are expressed in the hypothalamus making them possible candidates for compensatory mechanisms. Of these *fgf*s the only one being upregulated in homozygous *fgf3*^*t24152*^ mutants and passing the defined fold change threshold both at 3 and 7 dpf was *fgf3* itself, arguing for a self-compensatory role. Further, *fgf11b* was upregulated in mutants, but only at 7 dpf. *fgf8a* also showed a mild upregulation, but did not pass the defined threshold. *fgf8a* transcripts have been detected by *in situ* hybridisation in the posterior hypothalamus, but are spatially more restricted compared to *fgf3* (Reifers et al., 1998). The differential expression of downstream Fgf-signalling and feedback regulator genes was moderate in the *fgf3*^*t24152*^ mutants, but a few of them were downregulated at 7 dpf. Among these were three *dusp* genes (*dusp1, 2* and *5*), which act as negative feedback regulators of mitogen-activated protein kinases (Caunt and Keyse, 2013; Ornitz and Itoh, 2015; Znosko et al., 2010). Also *dusp6* showed a mild downregulation, but did not fulfil our selection criteria. Taken together, these findings suggest that a loss of Fgf3 activity in the *fgf3*^*t24152*^ mutant is compensated for, but only mildly, on multiple levels by both self-compensation and by other Fgf ligands, as well as by alterations in the downstream signalling.

In addition to Fgf-signalling, other signalling pathways and gene regulatory networks are active in the zebrafish developing posterior hypothalamus. For instance, *lef1*, a direct mediator of Wnt-signalling, is transcribed there, and is required for the expression of proneural and neuronal genes (Lee et al., 2006). A likely Wnt candidate in this context is Wnt8b (Lee et al., 2006). Supporting a role for Wnt-signalling in hypothalamic serotonergic neurogenesis, a subset of 5-HT immuno-reactive hypothalamic cells express Wnt-activity reporters (Lee et al., 2006; Wang et al., 2009, 2012). Further, functional studies have shown that hypothalamic Wnt-responsive cells are immature cells contributing to GABAergic and serotonergic populations (Wang et al., 2012). Further, functional impairment of *fezf2* in zebrafish results in fewer serotonergic, dopaminergic and oxytocinergic cells (Blechman et al., 2007; Guo et al., 1999; Jeong et al., 2006; Levkowitz et al., 2003; Rink and Guo, 2004), and at least for the dopaminergic populations this seems to involve a Fezf2-dependant regulation of the transcription factors Neurogenin 1 and Orthopedia (Jeong et al., 2006; Shimizu and Hibi, 2009; Blechman et al., 2007; Ryu et al., 2007). If, and to which extent, these signalling pathways interact with Fgf-signalling to promote the generation of monoaminergic CSF-c cells remains a subject of investigation.

## Acknowledgements

We are grateful to M. Schartl for generously providing access to fish facility and lab infrastructure. Furthermore, we are thankful to C. Drepper, S. Meierjohann and C. Stigloher for discussions and input on the project. The *fgf3*^*t24152*^ mutant was kindly provided by the M. Hammerschmidt lab, and plasmids by the L. Bally-Cuif lab.

## Data availability

Datasets will be made publicly available at the time of publication.

## Figure legends

**Suppl. Fig. 1.**
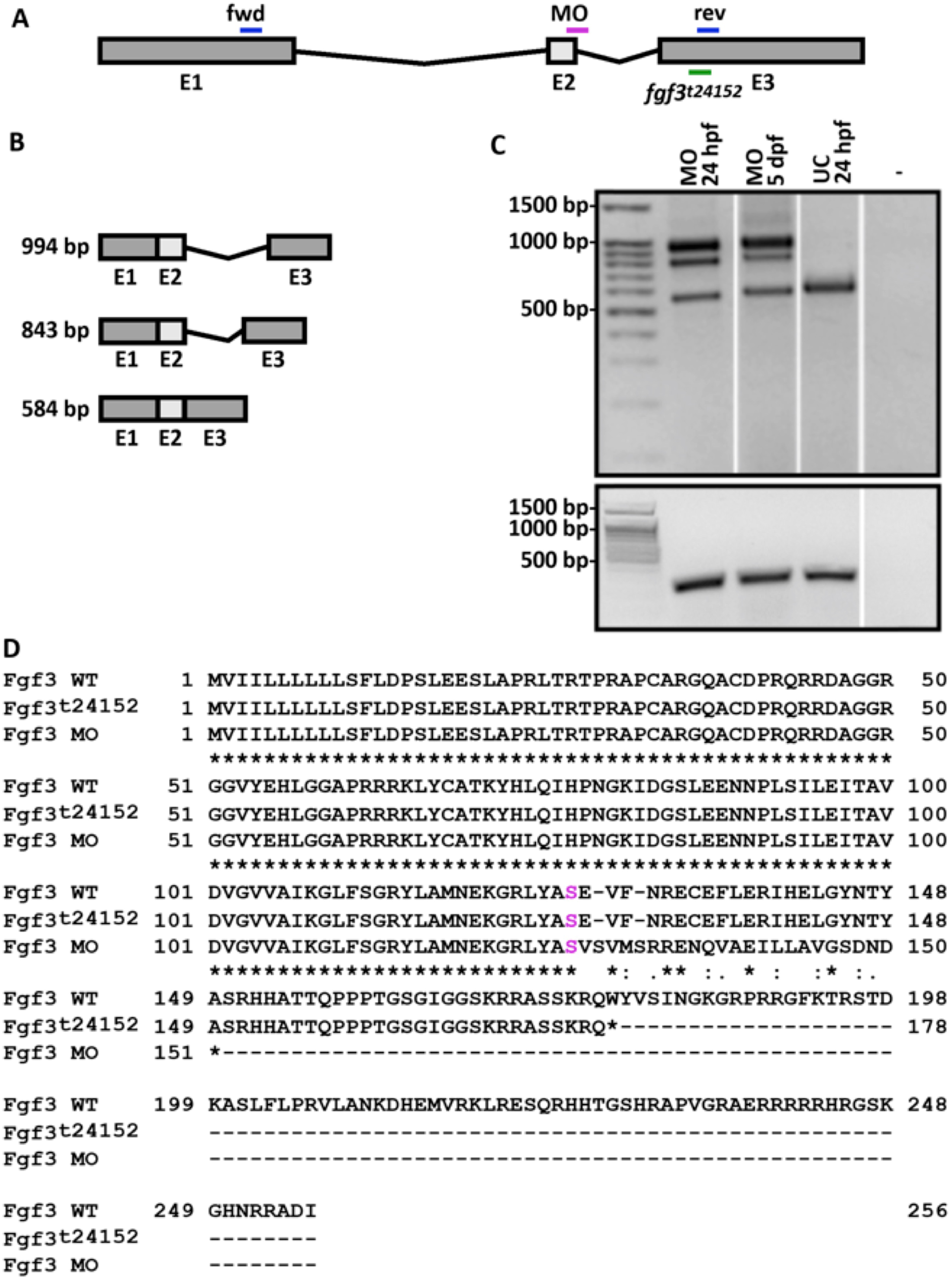
*fgf3* morpholino (MO) knock-down efficiency and amino acid sequence alignment of wildtype (WT), mutant (*fgf3*^*t24152*^) and morphant Fgf3. **(A)** Scheme of *fgf3* gene and locations of splice MO target site (magenta), as well as forward (fwd) and reverse (rev) primers (blue) used for detection of morphant splice forms, and point mutation *fgf3*^*t24152*^ (green). **(B)** Schematic representation of splice products detected by RT-PCR after MO injections. **(C)** RT-PCR results of morphants at 24 hpf and 5 dpf, uninjected control (UC) siblings at 24 hpf and water control (-) for *fgf3* (upper gel) and *β-actin* (lower gel). Additional RT-PCR products in *fgf3* morphants at 843 and 994 bp corresponded to a partial intron 2 inclusion due to a cryptic splice site or a complete inclusion of intron 2, respectively. **(D)** Amino acid sequence alignment of WT, *fgf3*^*t24152*^ mutant and morphant Fgf3. The *fgf3*^*t24152*^ mutation results in a premature stop after amino acid 177 (Q), thus, 69.1% of the WT protein sequence remains. After MO application, both splice forms of *fgf3* generate a nonsense sequence after amino acid 127 (S, magenta) and a stop after amino acid 150 (D) thus 49.6% of the WT protein sequence is intact.

**Suppl. Fig. 2.**
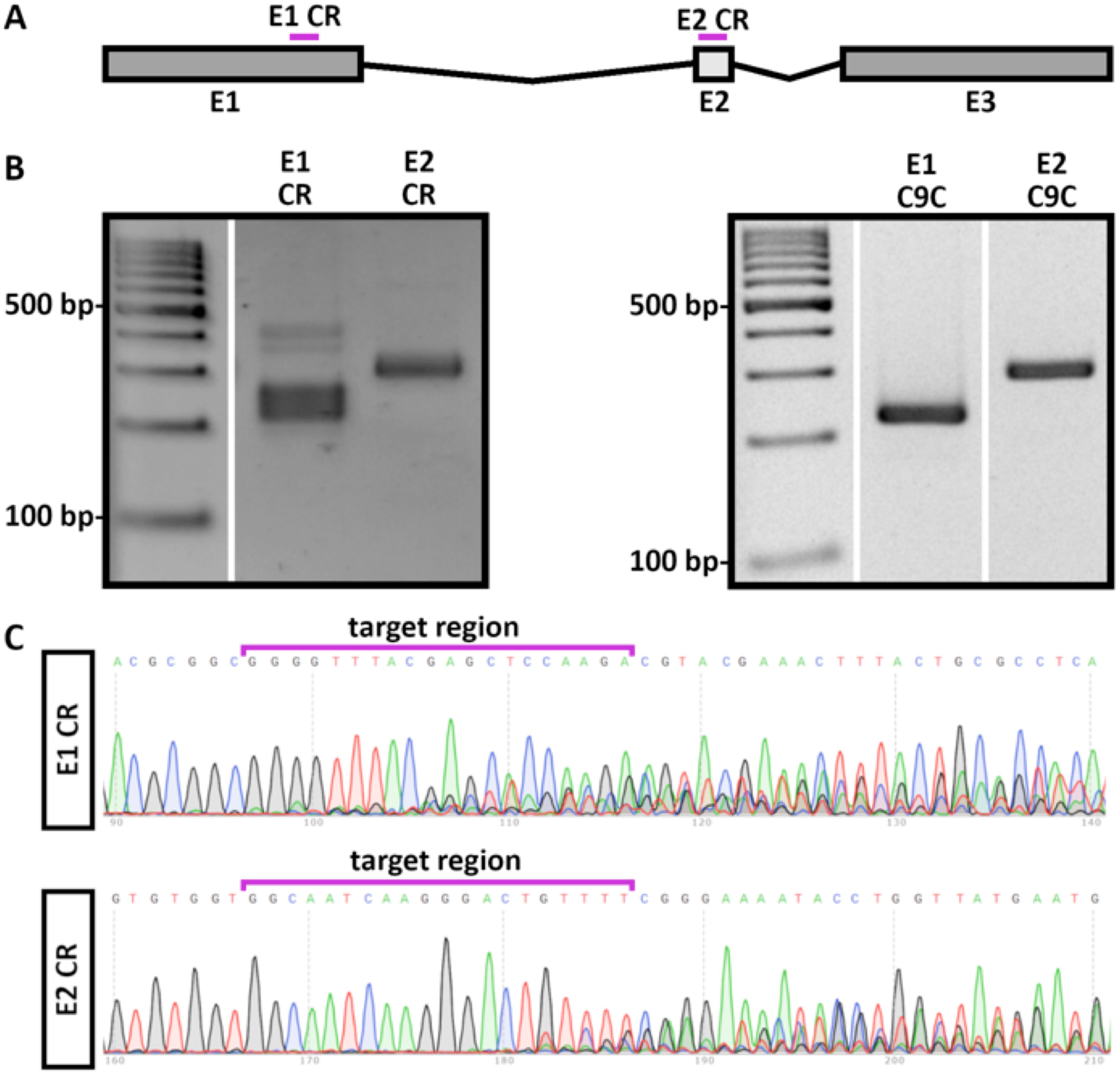
Validation of *fgf3* CRISPR/Cas9. **(A)** Scheme of *fgf3* gene with location of gRNA target region in exon 1 (E1 CR) and exon 2 (E2 CR) highlighted in magenta. **(B)** Examples of genotyping results from one 72 hpf embryo injected with both gRNAs together with Cas9 (left gel) and one 72 hpf control embryo injected with Cas9 only (C9C) (right gel). Note multiple bands for E1 CR and E2 CR gRNA in CRISPR/Cas9 injected embryo. The control embryo shows single wildtype bands (E1 C9C: 224 bp, E2 C9C: 289 bp) as result of PCR amplification of E1 and E2 target region. **(C)** DNA sequencing traces of genotyping PCR of CRISPR/Cas9 injected embryo (same specimen as for left gel in **B)** for E1 and E2 gRNA target regions revealed multiple traces due to indel mutations.

**Suppl. Fig. 3.**
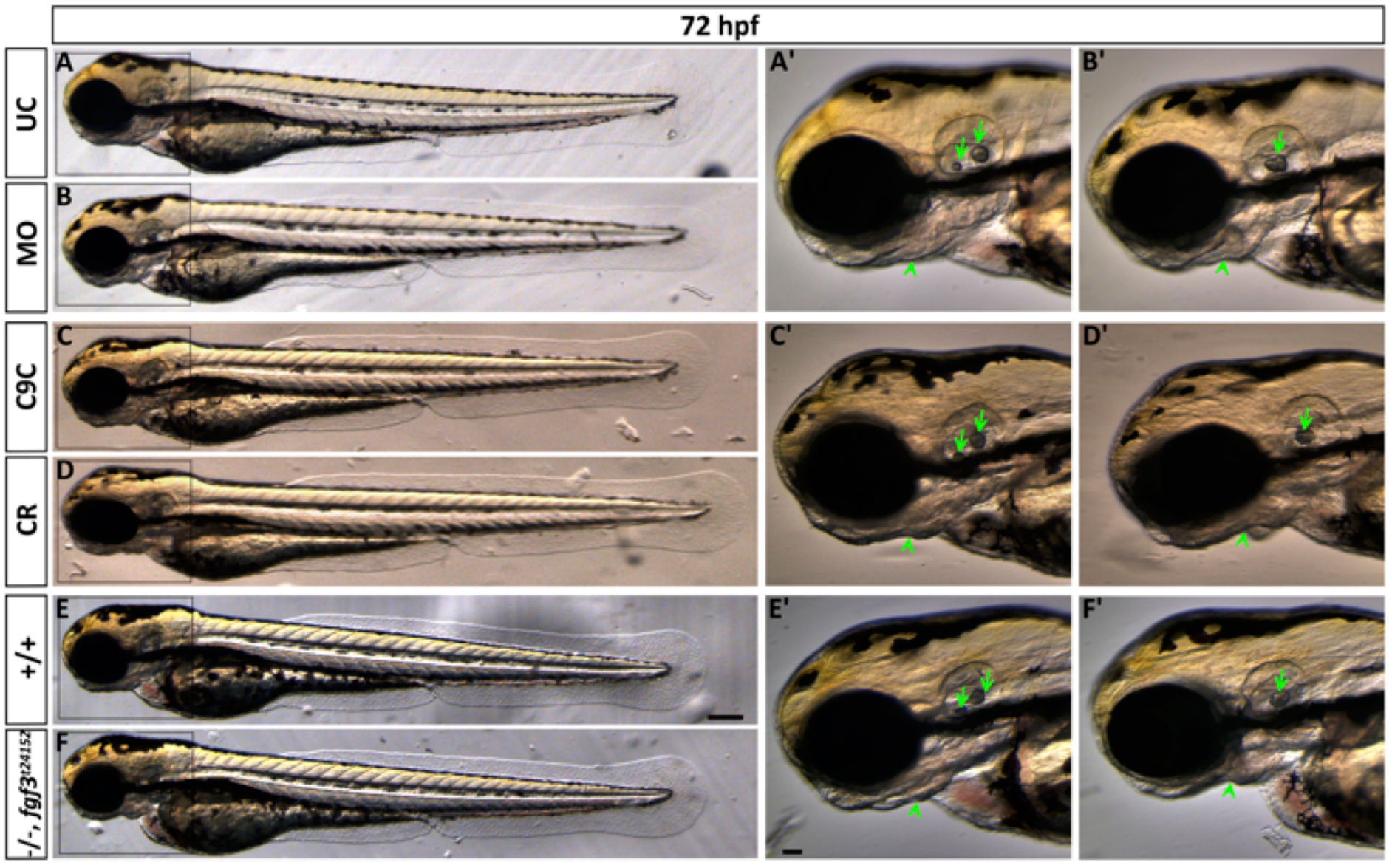
Live images showing morphology of 72 hpf embryos after *fgf3* impairment with characteristic ear and craniofacial malformations. (**A, B**) Uninjected control (UC) and *fgf3* morpholino injected (MO) siblings. (**C, D**) Cas9 injected control (C9C) and *fgf3* CRISPR/Cas9 injected (CR) siblings. (**E, F**) Wildtype (+/+) and homozygous *fgf3*^*t24152*^ mutant (-/-) siblings. Boxes indicate magnified area shown in A’-F’. After *fgf3* impairment the two otoliths fuse (arrow) and the lower jaw bones are malformed (arrow heads). Lateral views, anterior to the left. Scale bar in E, 100 µm; in E’, 50 µm.

**Suppl. Fig. 4.**
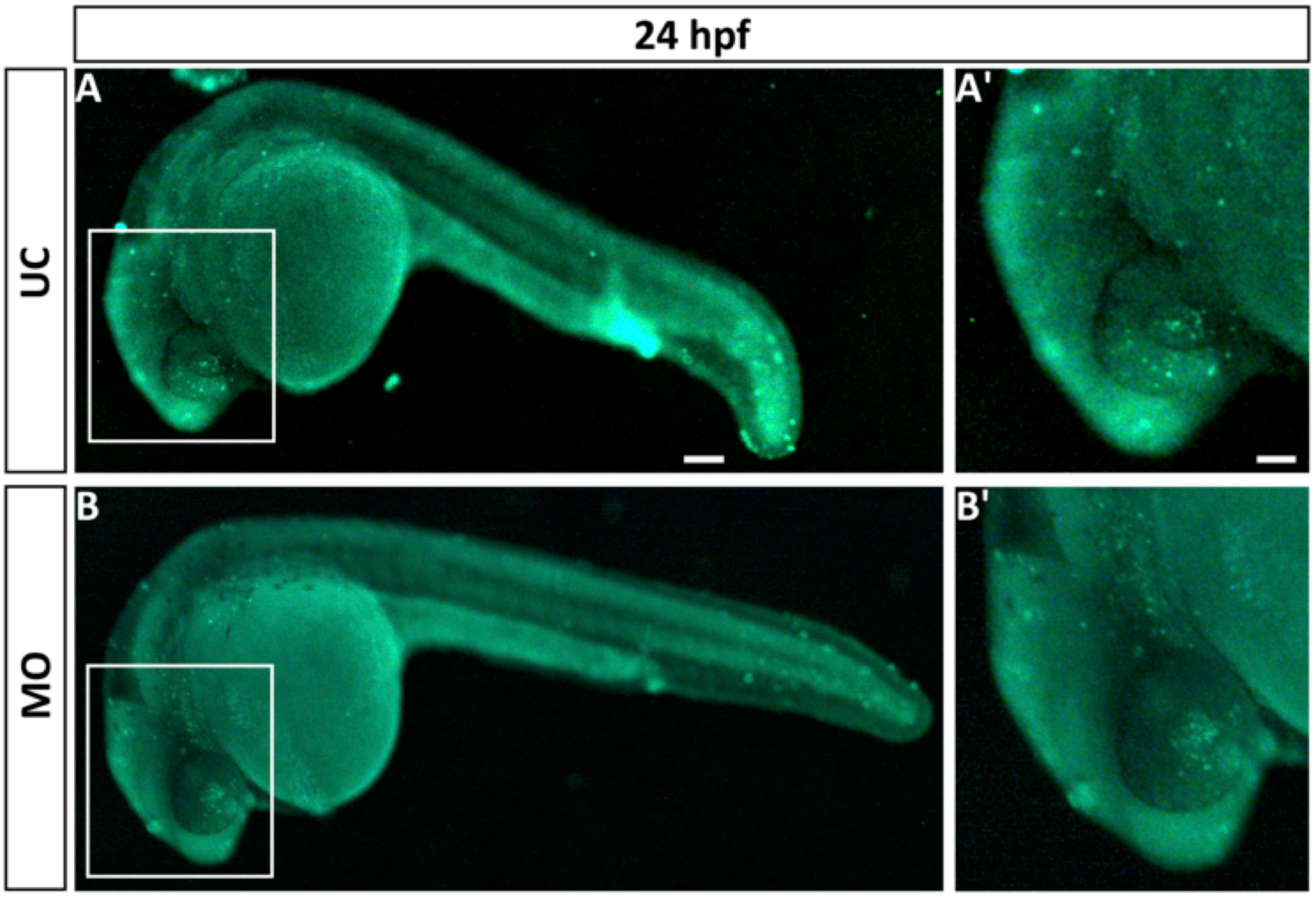
Live fluorescence pictures showing cell death in *fgf3* morphants at 24 hpf. **(A, B)** Acridine orange staining of uninjected control (UC) and morpholino injected (MO) siblings at 24 hpf. Boxes indicate magnified area shown in A’ and B’. Lateral views, anterior to the left. Scale bar in A, 100 µm; in A’, 50 µm.

**Suppl. Fig. 5.**
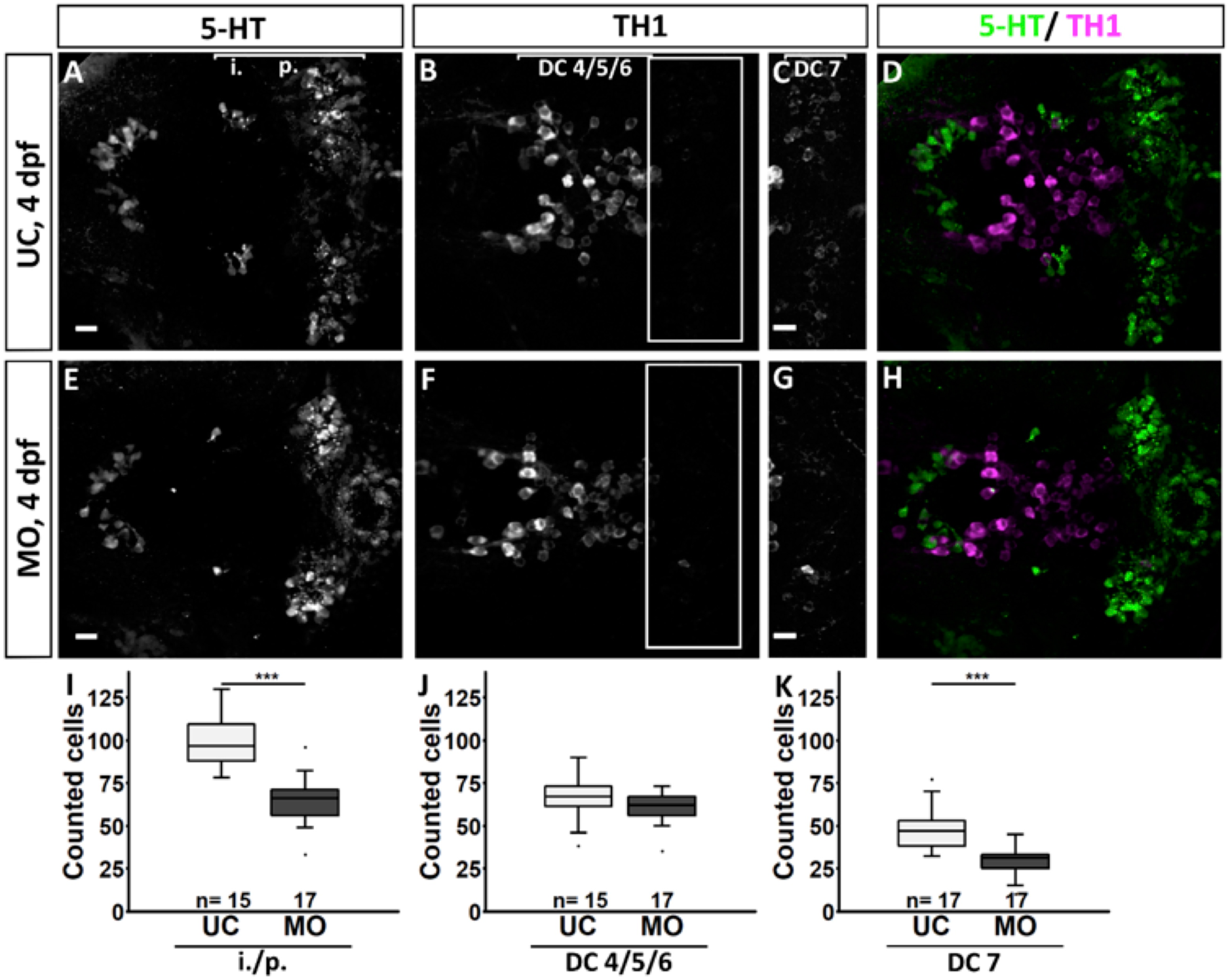
Quantification of the number of serotonergic cells in the intermediate (i.)/posterior (p.) clusters and of dopaminergic cells in the DC 4/5/6 and DC 7 clusters in the hypothalamus of *fgf3* morphants at 4 dpf. **(A-H)** Confocal maximum intensity projections from uninjected control (UC) and morpholino injected (MO) siblings immuno stained for 5-HT (green) and TH1 (magenta) shown as single channels (**A-C, E-G**) and merged (**D, H**). **C** and **G** show boxed areas in **B** and **F**, respectively, with adjusted brightness and contrast to reveal the faint TH1 immunoreactive cells of the DC 7 cluster. Ventral views, anterior to the left. Scale bars = 10 µm. **(I-K)** Quantifications of 5-HT and TH1 positive cells in control and morphant siblings. The number of serotonergic cells was counted in the i./p. clusters as indicated by the line in **A**. The number of dopaminergic cells was counted in the DC 4/5/6 and DC 7 clusters as indicated by the lines in **B** and **C**, respectively. Boxplots show median, 25-75% percentile, min/max whiskers and outliers depicted as dots. n = number of analysed individuals.

**Suppl. Table 1.**
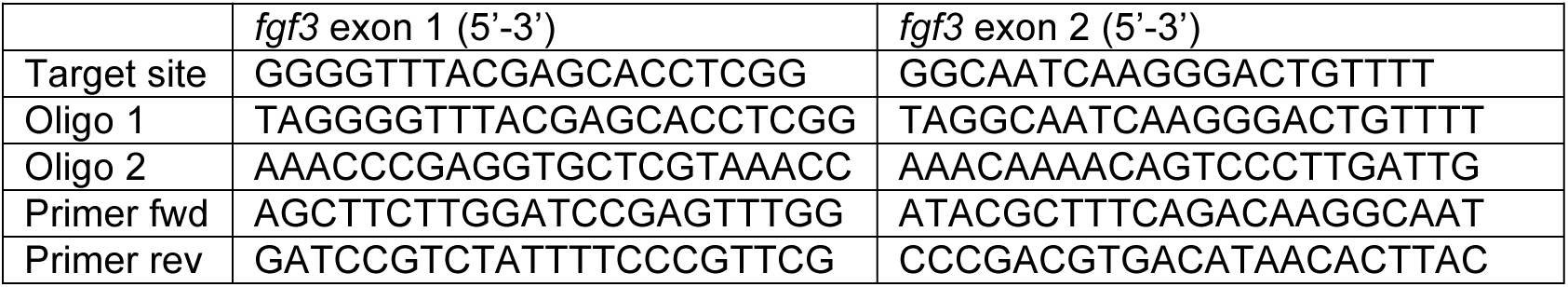
Target sites, oligos and PCR primers for *fgf3* CRISPR/Cas9 approach.

**Suppl. Table 2.** RNA probes used for whole mount RNA *in situ* hybridisation.

| Gene name | Gene symbol | Reference |
| --- | --- | --- |
| *arginine vasopressin* | *avp* | Eaton et al., 2008 |
| *cortistatin* | *cort* | Devos et al., 2002 |
| *ets variant 5b* | *etv5b* | Münchberg et al., 1999 |
| *fibroblast growth factor 3* | *fgf3* | Kiefer et al., 1996 |
| *NK2 homeobox 4b* | *nkx2.4b* | Rohr & Concha, 2000 |
| *oxytocin* | *oxt* | Unger & Glasgow, 2003 |

**Suppl. Table 3.**
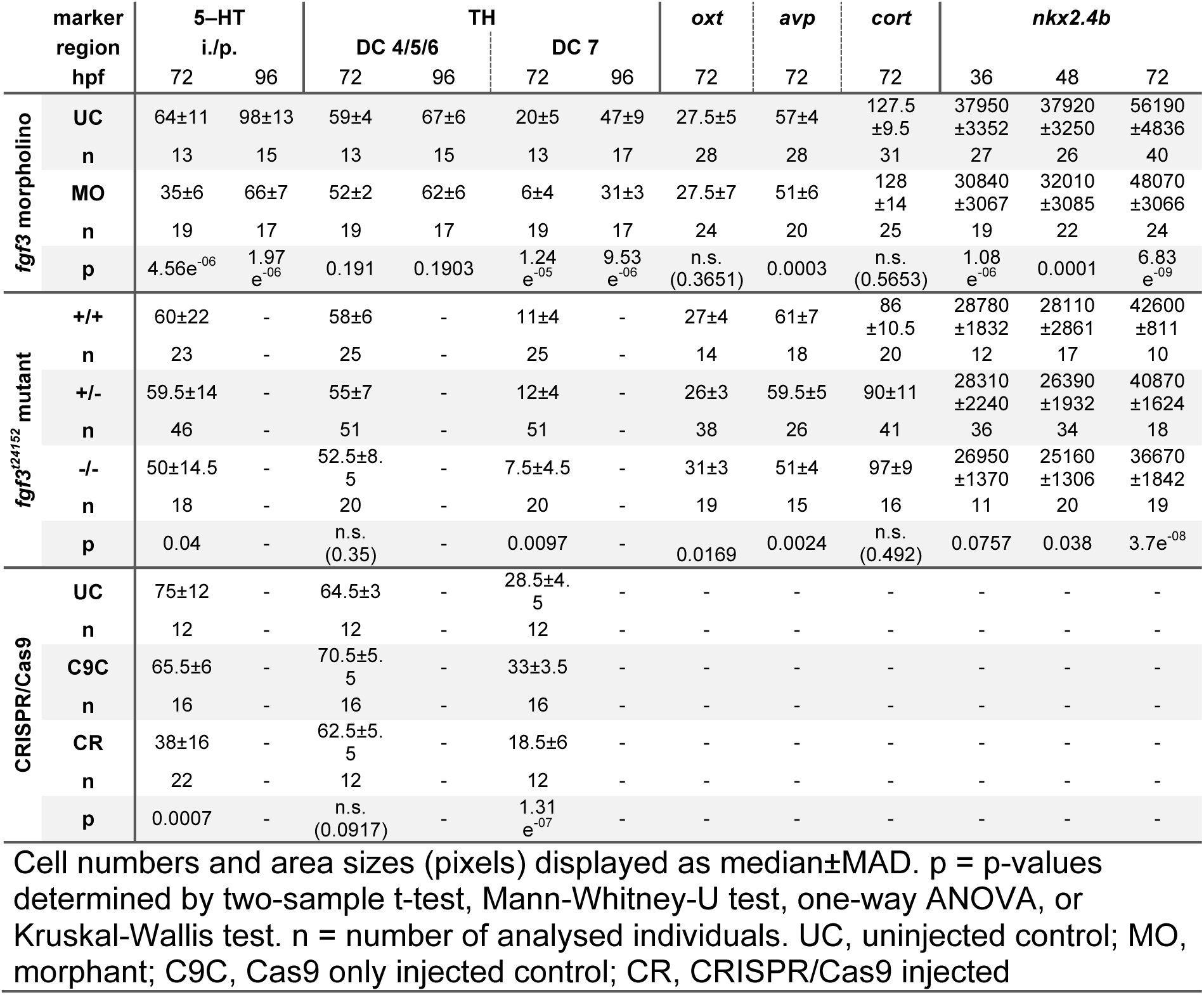
Overview of counting of monoaminergic and neuroendocrine cells, and of measurements of the *nkx2.4b* expressing hypothalamic area in embryos after *fgf3* impairment.

**Suppl. Table 4:**
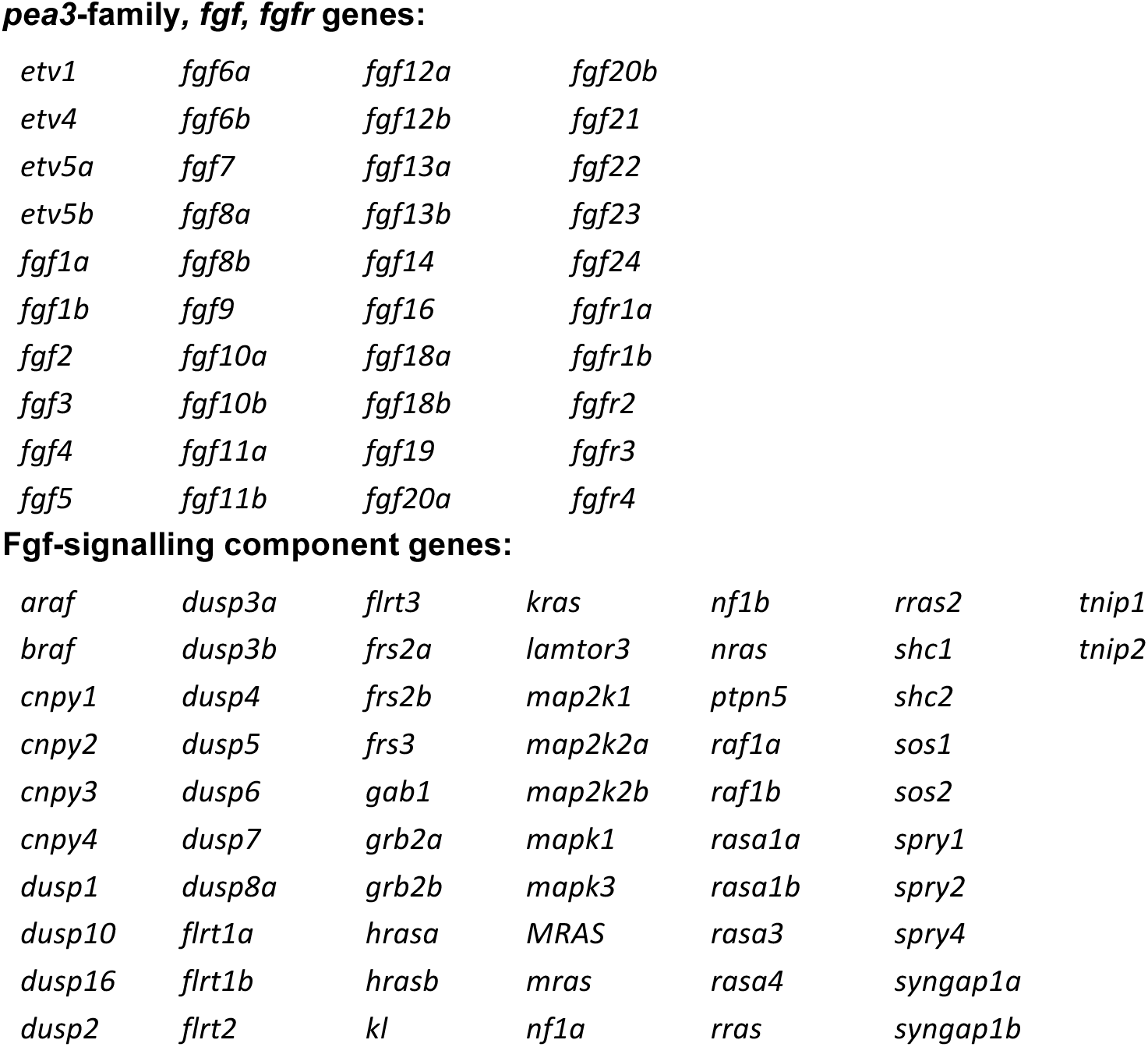
List of Fgf-signalling genes selected for expression analysis in the hypothalamus by RNA sequencing.

***pea3*-family, *fgf*, *fgfr* genes:**
|  |  |  |  |
| --- | --- | --- | --- |
| *etv1* | *fgf6a* | *fgf12a* | *fgf20b* |
| *etv4* | *fgf6b* | *fgf12b* | *fgf21* |
| *etv5a* | *fgf7* | *fgf13a* | *fgf22* |
| *etv5b* | *fgf8a* | *fgf13b* | *fgf23* |
| *fgf1a* | *fgf8b* | *fgf14* | *fgf24* |
| *fgf1b* | *fgf9* | *fgf16* | *fgfr1a* |
| *fgf2* | *fgf10a* | *fgf18a* | *fgfr1b* |
| *fgf3* | *fgf10b* | *fgf18b* | *fgfr2* |
| *fgf4* | *fgf11a* | *fgf19* | *fgfr3* |
| *fgf5* | *fgf11b* | *fgf20a* | *fgfr4* |

**Fgf-signalling component genes:**
|  |  |  |  |  |  |  |
| --- | --- | --- | --- | --- | --- | --- |
| *araf* | *dusp3a* | *flrt3* | *kras* | *nf1b* | *rras2* | *tnip1* |
| *braf* | *dusp3b* | *frs2a* | *lamtor3* | *nras* | *shc1* | *tnip2* |
| *cnpy1* | *dusp4* | *frs2b* | *map2k1* | *ptpn5* | *shc2* |  |
| *cnpy2* | *dusp5* | *frs3* | *map2k2a* | *raf1a* | *sos1* |  |
| *cnpy3* | *dusp6* | *gab1* | *map2k2b* | *raf1b* | *sos2* |  |
| *cnpy4* | *dusp7* | *grb2a* | *mapk1* | *rasa1a* | *spry1* |  |
| *dusp1* | *dusp8a* | *grb2b* | *mapk3* | *rasa1b* | *spry2* |  |
| *dusp10* | *flrt1a* | *hrasa* | *MRAS* | *rasa3* | *spry4* |  |
| *dusp16* | *flrt1b* | *hrasb* | *mras* | *rasa4* | *syngap1a* |  |
| *dusp2* | *flrt2* | *kl* | *nf1a* | *rras* | *syngap1b* |  |

Author contributions: Designed the study (IR, KPL, CL); wrote the manuscript (IR, CL); edited the manuscript (all); performed the experiments (IR, JJ, SK, JK, CL)

